# A mathematical approach to differentiate spontaneous and induced evolution to drug resistance during cancer treatment

**DOI:** 10.1101/235150

**Authors:** James M. Greene, Jana L. Gevertz, Eduardo D. Sontag

**Affiliations:** Department of Mathematics and Center for Quantitative Biology, Rutgers University; Department of Mathematics and Statistics, The College of New Jersey; Department of Bioengineering and Department of Electrical and Computer Engineering, Northeastern University; Harvard Program in Therapeutic Science, Harvard Medical School

**Keywords:** Drug resistance, Chemotherapy, Phenotype, Identifiability, Control theory

## Abstract

Drug resistance is a major impediment to the success of cancer treatment. Resistance is typically thought to arise through random genetic mutations, after which mutated cells expand via Darwinian selection. However, recent experimental evidence suggests that the progression to drug resistance need not occur randomly, but instead may be induced by the treatment itself, through either genetic changes or epigenetic alterations. This relatively novel notion of resistance complicates the already challenging task of designing effective treatment protocols. To better understand resistance, we have developed a mathematical modeling framework that incorporates both spontaneous and drug-induced resistance. Our model demonstrates that the ability of a drug to induce resistance can result in qualitatively different responses to the same drug dose and delivery schedule. We have also proven that the induction parameter in our model is theoretically identifiable, and proposed an *in vitro* protocol which could be used to determine a treatment’s propensity to induce resistance.

## Introduction

Tumor resistance to chemotherapy and targeted drugs is a major cause of treatment failure. Both molecular and microenvironmental factors have been implicated in the development of drug resistance [33]. As an example of molecular resistance, the upregulation of drug efflux transporters can prevent sufficiently high intracellular drug accumulation, limiting treatment efficacy [30]. Other molecular causes of drug resistance include modification of drug targets, enhanced DNA damage repair mechanisms, dysregulation of apoptotic pathways, and the presence of cancer stem cells [30, 17, 33, 79, 82]. The irregular tumor vasculature which results in inconsistent drug distribution and hypoxia is an example of a microenvironmental factor that impacts drug resistance [77]. Other characteristics of the tumor microenvironment that influence drug resistance include regions of acidity, immune cell infiltration and activation, and the tumor stroma [26, 77, 55, 15, 33, 54]. Experimental and clinical research continues to shed light on the multitude of factors that contribute to cancer drug resistance. Mathematical modeling studies have also been used to explore both broad and detailed aspects of cancer drug resistance, as reviewed in [42, 7, 24].

Resistance to cancer drugs can be classified as either pre-existing or acquired [33]. Preexisting (intrinsic) drug resistance describes the case in which a tumor contains a subpopulation of drug resistant cells at the initiation of treatment, making the therapy (eventually) ineffective due to resistant cell selection [33]. As examples, pre-existing BCR-ABL kinase domain mutations confer resistance to the tyrosine kinase inhibitor imatinib in chronic myeloid leukemia patients [66, 35], and pre-existing MEK1 mutations confer resistance to BRAF inhibitors in melanoma patients [8]. Many mathematical models have considered how the presence of such pre-existing resistant cells impact cancer progression and treatment [36, 38, 57, 39, 22, 23, 71, 16, 59, 62, 4, 21, 43, 3, 31, 81, 28, 48, 56, 58, 78, 69, 9, 10].

On the other hand, acquired drug resistance broadly describes the case in which drug resistance develops during the course of therapy from a population of cells that were initially drug sensitive [33]. The term “acquired resistance” is really an umbrella term for two distinct phenomena, which complicates the study of acquired resistance. On the one hand there is resistance that is *spontaneously, or randomly, acquired* during the course of treatment, be it due to random genetic mutations or stochastic non-genetic phenotype switching [65]. This spontaneous form of acquired resistance has been considered in many mathematical models [14, 38, 46, 57, 39, 22, 23, 44, 4, 21, 32, 43, 51, 31, 81, 25, 45, 50, 56, 10]. On the other hand, drug resistance can be *induced (caused)* by the drug itself [61, 60, 70, 65, 64].

The question of whether resistance is an induced phenomenon or the result of random events was first famously studied by Luria and Delbrück in the context of bacterial (*Escherichia coli*) resistance to a virus (T1 phage) [52]. In particular, Luria and Delbrück hypothesized that if selective pressures imposed by the presence of the virus *induce* bacterial evolution, then the number of resistant colonies that formed in their plated experiments should be Poisson distributed, and hence have an approximately equal mean and variance. What Luria and Delbrück found instead was that the number of resistant bacteria on each plate varied drastically, with the variance being significantly larger than the mean. As a result, they concluded that the bacterial mutations are spontaneous, and not induced by the presence of the virus [52].

In the case of cancer, there is strong evidence that at least some drugs have the ability to induce resistance, as genomic mutations can be caused by cytotoxic cancer chemotherapeutics [74, 73]. For instance, nitrogen mustards can induce base substitutions and chromosomal rearrangements, topoisomerase II inhibitors can induce chromosomal translocations, and antimetabolites can induce double stranded breaks and chromosomal aberrations [74]. Such drug-induced genomic alterations would generally be non-reversible. Drug resistance can also be induced at the epigenetic level [61, 65, 68]. As one example, the expression of multidrug resistance 1 (MDR1), an ABC-family membrane pump that mediates the active efflux of drug, can be induced during treatment [33, 65]. In another recent example, the addition of a chemotherapeutic agent is shown to induce, through a multistage process, epigenetic reprogramming in patient-derived melanoma cells [68]. Resistance developed in this way can occur quite rapidly, and can often be reversed [70, 65, 34].

Despite these known examples of drug-induced resistance, differentiating between drug-selected and drug-induced resistance is nontrivial. What appears to be drug-induced acquired resistance may simply be the rapid selection of a very small number of pre-existing resistant cells, or the selection of cells that spontaneously acquired resistance [44, 65]. In pioneering work by Pisco and colleagues, the relative contribution of resistant cell selection versus drug-induced resistance was assessed in an experimental system involving HL60 leukemic cells treated with the chemotherapeutic agent vincristine [65]. After 1-2 days of treatment, the expression of MDR1 was shown to be predominantly mediated by cell-individual induction of MDR1 expression, and not by the selection of MDR1-expressing cells [65, 64]. In particular, these cancer cells exploit their heritable, non-genetic phenotypic plasticity (by which one genotype can map onto multiple stable phenotypes) to change their gene expression to a (temporarily) more resistant state in response to treatment-related stress [65, 64].

Although there is a wealth of mathematical research addressing cancer drug resistance, relatively few models have considered drug-induced resistance. Of the models of drug-induced resistance that have been developed, many do not explicitly account for the presence of the drug. Instead, it is assumed that these models apply only under treatment [65, 29, 11, 1, 19], with the effects of the drug being implicitly captured in the model terms. Since these models of resistance-induction are dose-independent, they are unable to capture the effects that altering the drug dose has on resistance formation. To our knowledge, there have been less than a handful of mathematical models developed in which resistance is induced by a drug in a dose-dependent fashion [13, 28, 48]. In [28] and follow-up work in [69, 63], the duration and intensity of drug exposure determines the resistance level of each cancer cell. This model allows for a continuum of resistant phenotypes, but is very computationally complex as it is a hybrid discrete-continuous, stochastic spatial model. While interesting features about the relationship between induced resistance and the microenvironment have been deduced from this model, its complexity does not allow for general conclusions to be drawn about dose-dependent resistance-induction.

Another class of models which addresses drug-induced resistance is that in [13]. These models are distinct in that they are motivated by *in vitro* experiments in which a cancer drug transiently induces a reversible resistant phenotypic state [70]. The individual-based and integro-differential equation models developed consider rapidly proliferating drug sensitive cells, slowly proliferating drug resistant cells, and rapidly proliferating drug resistant cells. An advection term (with the speed depending on the drug levels) is used to model drug-induced adaptation of the cell proliferation level, and a diffusion term for both the level of cell proliferation and survival potential (response to drug) is used to model non-genetic phenotype instability [13]. Through these models, the contribution of nongenetic phenotype instability (both drug-induced and random), stress-induced adaptation, and selection can be quantified [13].

Finally, the work in [48] models the evolutionary dynamics of the tumor population as a multi-type non-homogeneous continuous-time birth-death stochastic process. This model accounts for the ability of a targeted drug to alter the rate of resistant cell emergence in a dose-dependent manner. The authors’ specifically considered the case where the rate of mutation that gives rise to resistant cell: 1) increases as a function of drug concentration, 2) is independent of drug concentration, and 3) decreases with drug concentration. Interestingly, this model led to the conclusion that the optimal treatment strategy is independent of the relationship between the drug concentration and rate of resistance formation. In particular, they found resistance is optimally delayed using a low-dose continuous treatment strategy coupled with high-dose pulses [48].

As *in vitro* experiments have demonstrated that treatment response can be impacted by drug-induced resistance [70, 65], here we seek to understand this phenomenon further using mathematical modeling. The initial mathematical model that we developed, and that will be analyzed herein, is a system of two ordinary differential equations with a single control representing the drug. We intentionally chose a minimal model that would be amenable to analysis, as compared to previously-developed models of drug-induced resistance which are significantly more complex [13, 28, 69, 63]. Despite the simplicity of the model, it incorporates both spontaneous and drug-induced resistance.

This manuscript is organized as follows. We begin by introducing a mathematical model to describe the evolution of drug resistance during treatment with a theoretical resistance-inducing (and non-inducing) drug. We use this mathematical model to explore the role that the drug’s resistance induction rate has on treatment dynamics. We demonstrate that the induction rate of a theoretical cancer drug could have a nontrivial impact on the qualitative responses to a given treatment strategy, including the tumor composition and the time horizon of tumor control. In our model, for a *resistance-preserving* drug (i.e. a drug which does not induce resistance), better tumor control is achieved using a constant therapeutic protocol as compared to a pulsed one. On the other hand, in the case of a resistance-inducing drug, pulsed therapy prolongs tumor control longer than does constant therapy due to sensitive/resistant cell competitive inhibition. Once the importance of induced resistance has been established, we demonstrate that all parameters in our mathematical model are identifiable, meaning that it is theoretically possible to determine the rate at which drug resistance is induced for a given treatment protocol. Since this theoretical result does not directly lend itself to an experimental approach for quantifying a drug’s ability to induce resistance, we also describe a potential *in vitro* experiment for approximating this ability for constant therapies. We end with some concluding remarks and a discussion of potential extensions of our analysis, such as a model that differentiates between reversible and non-reversible forms of resistance.

## Materials and Methods

In this section, we introduce a general modeling framework to describe the evolution of drug resistance during treatment. Our model captures the fact that resistance can result from random events, or can be induced by the treatment itself. Random events that can confer drug resistance include genetic alterations (e.g. point mutations or gene amplification) and phenotype-switching [65]. These spontaneous events can occur either prior to or during treatment. Drug-induced resistance is resistance specifically activated by the drug, and as such, depends on the effective dosage encountered by a cell. Such a formulation allows us to distinguish the contributions of both drug-dependent and drug-independent mechanisms, as well as any dependence on preexisting (i.e. prior to treatment) resistant populations.

We consider the tumor to be composed of two types of cells: sensitive (S) and resistant (R). Sensitive (or wild-type) cells are fully susceptible to treatment, while treatment affects resistant cells to a lesser degree. To analyze the role of both random and drug-induced resistance, we utilize a system of two ordinary differential equations (ODEs) to describe the dynamics between the S and R subpopulations:

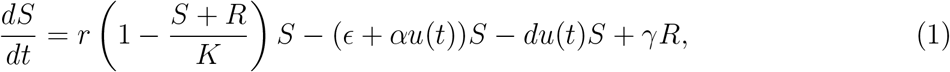

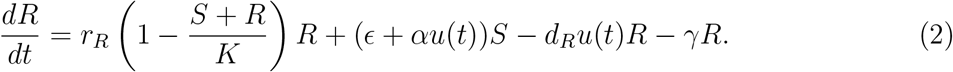

All parameters are non-negative. In the absence of treatment, we assume that the tumor grows logistically, with each population contributing equally to competitive inhibition. Phenotypes *S* and *R* each possess individual intrinsic growth rates, and we make the assumption in the remainder of the work that 0 < *r_R_* < *r*. This simply states that resistant cells grow slower than non-resistant cells, an assumption supported by experimental evidence [47, 67, 6].

The transition to resistance can be described with a net term of the form *ϵS* + *αu*(*t*)*S*. Mathematically, the drug-induced term *αu*(*t*)*S*, where *u*(*t*) is the effective applied drug dosage at time *t*, describes the effect of treatment on promoting the resistant phenotype. As an example, this term could represent the induced over-expression of the P-glycoprotein gene, a well-known mediator of multi-drug resistance, by the application of chemotherapy [33, 75].

The spontaneous evolution of resistance is captured in the *ϵS* term, which permits resistance to develop even in the absence of treatment. Note that *ϵ* is generally considered small [49], although recent experimental evidence into error prone DNA polymerases suggests that cancer cells may have increased mutation rates due to the over-expression of such polymerases [40, 53, 41]. For example, in [40], mutation rates due to such polymerases are characterized by probabilities as high as 7.5 × 10^-1^ per base substitution, and it is known that many point mutations in cancer arise from these DNA polymerases [41]. For this work we adopt the notion that random point mutations leading to drug resistance are rare, and that drug-induced resistance occurs on much quicker time scales [65]. Therefore we will assume that *α* > *ϵ* in our analysis of eqns. (1)-(2).

We model the effects of treatment by assuming the log-kill hypothesis [76], which states that a given dose of chemotherapy eliminates the same fraction of tumor cells, regardless of tumor size. We allow for each cellular compartment to have a different drug-induced death rate (*d*, *d_R_*); however, to accurately describe resistance it is required that 0 ≤ *d_R_* < *d*. Our analysis presented herein will be under the simplest assumption that the drug is completely ineffective against resistant cells, so that *d_R_* = 0.

The last term in the equations, *γR*, represents the re-sensitization of cancer cells to the drug. In the case of non-reversible resistance, *γ* = 0 and otherwise *γ* > 0. Our subsequent analysis will be done under the assumption of non-reversible resistance. For a discussion of the effect of reversibility on the presented model, see Supporting Information (SI) Section B.

Finally, we note that the effective drug concentration *u*(*t*) can be thought of as a control input. For simplicity, in this work we assume that it is directly proportional to the applied drug concentration. However, pharmacodynamic/pharmacokinetic considerations could be incorporated to more accurately describe the uptake/evolution of the drug *in vivo* or *in vitro*; for example, as in [2, 80, 20].

To understand the above system of drug resistance evolution, we reduce the number of parameters via non-dimensionalization. Rescaling *S* and *R* by their (joint) carrying capacity *K*, and time *t* by the sensitive cell growth rate *r*,

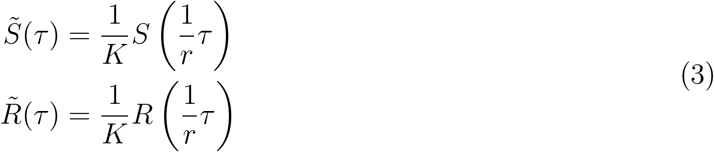

system (1)-(2) (with *γ* = *d_R_*= 0) can be written in the form

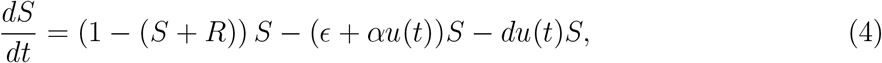

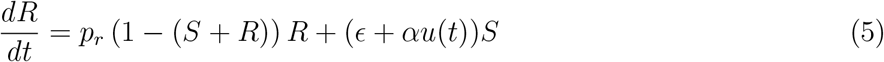

For convenience, we have relabeled *S*, *R*, and *t* to coincide with the non-dimensionalization, so that the parameters *ϵ*, *α*, and *d* must be scaled accordingly (by 1/*r*). As *r_r_* was assumed to satisfy 0 ≤ *r_R_* < *r*, the relative resistant population growth rate *p_r_* satisfies 0 ≤ *p_r_* < 1.

One can show (see SI Section A) that asymptotically, under any treatment regime *u*(*t*) ≥ 0, the entire population will become resistant:

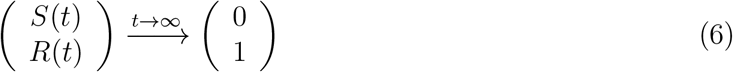

However, tumor control is still possible, where one can combine therapeutic efficacy and clonal competition to influence transient dynamics and possibly prolong patient life. Indeed, the modality of adaptive therapy has shown promise in utilizing real-time patient data to inform therapeutic modulation aimed at increasing quality of life and survival times [27]. This work will focus on such dynamics and controls.

## Results/Discussion

### Effect of Induction on Treatment Efficacy

We investigate the role of a drug’s induction capability (parameter α in (4)-(5)) on treatment dynamics. Specifically, the value ofα may have a substantial impact on the relative success of two standard therapy protocols: constant dosage and periodic pulsing.

#### Treatment Protocol

To quantify the effects of induced resistance, a treatment protocol must be specified. We adopt a clinical perspective over the course of the disease, which is summarized in Figure 1. We assume that the disease is initiated by a small number of wild-type cells:

**Figure 1:**
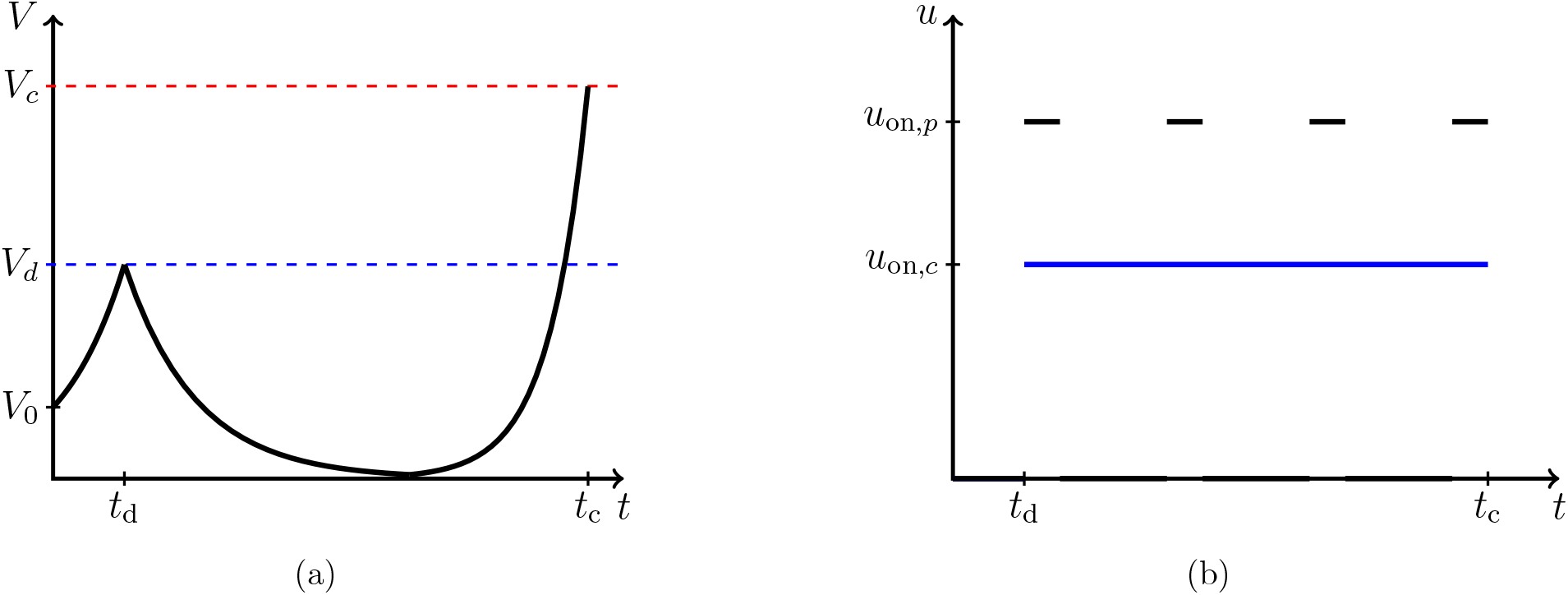
Schematic of tumor dynamics under two treatment regimes. (a) Tumor volume V in response to treatment initiated at time *t_d_*. Cancer population arises from a small sensitive population at time *t* = 0, upon which the tumor grows to detection at volume *V_d_*. Treatment is begun at *t_d_*, and continues until the tumor reaches a critical size *V_c_* (at a corresponding time *t_c_*), where treatment is considered to have failed. (b) Illustrative constant and pulsed treatments, both initiated at *t* = *t_d_*.

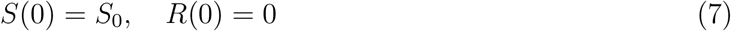

where 0 < *S*_0_ < 1. The tumor then progresses untreated until a specific volume *V_d_* is detected (or, for hematologic tumors, via appropriate blood markers), which utilizing existing nuclear imaging techniques corresponds to a tumor with diameter on the order of 10 mm [18]. The time to reach *V_d_* is denoted by *t_d_*, which in general depends on all parameters appearing in (4)-(5). Note that, assuming *ϵ* > 0, a non-zero resistant population will exist at the onset of treatment. Therapy, represented through *u*(*t*), is then applied, until the tumor reaches a critical size *V_c_*, which we equate with treatment failure. Since the (*S*, *R*) = (0,1) state is globally asymptotically stable in the first quadrant, *V_c_* < 1 is guaranteed to be obtained in finite time. Time until failure, *t_c_*, is then a measure of efficacy of the applied *u*(*t*).

Although a diverse set of inputs *u*(*t*) may be theoretically applied, presently we consider only strategies as illustrated in Figure 1(b). The blue curve in Figure 1(b) corresponds to a constant effective dosage *u_c_*(*t*) initiated at *t_d_* (administered approximately utilizing continuous infusion pumps and/or slow-release capsules), while the black curve represents a corresponding pulsed strategy *u_p_*(*t*), with fixed treatment windows and holidays. In general, we may allow for different magnitudes, *u*_on,*c*_ and *u*_on,*p*_, for constant and pulsed therapies respectively; for example to relate the total dosage applied per treatment cycle (*AUC* in clinical literature). However for simplicity we assume the same magnitude in the subsequent section (although see SI Section C for a normalized comparison). While these represent idealized therapies, such *u*(*t*) may form an accurate approximation to in *vitro* and/or in *vivo* kinetics. Note that the response *V*(*t*) illustrated in Figure 1(a) will not be identical (or even qualitatively similar) for both presented strategies, as will be demonstrated numerically.

#### Constant vs. Pulsed Therapy Comparison

To qualitatively demonstrate the role induced resistance plays in designing schedules for therapy, we consider two drugs with the same cytotoxic potential (i.e. same drug-induced death rate d), each possessing a distinct level of resistance induction (parameter α). A fundamental question is then whether there exist qualitative distinctions between treatment responses for each chemotherapy. More specifically, how does the survival time compare when scheduling is altered between constant therapy and pulsing? Does the optimal strategy (in this case, optimal across only two schedulings) change depending on the extent to which the drug induces resistance?

We fix two values of the induction parameter *α*:

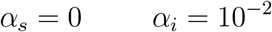

Recall that we are studying the non-dimensional model (4)-(5), so no units are specified. Parameter *α* = 0 corresponds to no therapy-induced resistance (henceforth denoted as *phenotype-preserving*), and therefore considering this case permits a comparison between the classical notion of random evolution towards resistance (*α* = 0) and drug-induced resistance (*α* > 0). For the remainder of the section, parameters are fixed as in Table 1. Importantly, all parameters *excluding α* are identical for each drug, enabling an unbiased comparison. Treatment magnitudes *u*_on,*c*_ and *u*_on,*p*_ are selected to be equal: *u*_on,*c*_ = *u*_on,*p*_ = *u*_on_.

**Table 1:** Parameters utilized in Figure 2

| Parameter | Biological Interpretation | Value (dimensionless) |
| --- | --- | --- |
| $S_0$ | Initial sensitive population | 0.01 |
| $R_0$ | Initial resistant population | 0 |
| $V_d$ | Detectable tumor volume | 0.1 |
| $V_c$ | Tumor volume defining treatment failure | 0.9 |
| $\epsilon$ | Background mutation rate | $10^{-6}$ |
| $d$ | Cytotoxicity of sensitive cells | 1 |
| $p_r$ | Resistant growth fraction | 0.2 |
| $u_{\text{on}}$ | Treatment magnitude, constant dose | 1.5 |
| $\Delta t_{\text{on}}$ | Pulsed treatment window | 1 |
| $\Delta t_{\text{off}}$ | Pulsed holiday length | 3 |

Note that selecting parameter *V_d_* =0.1 implies that the carrying capacity has a diameter of 100 mm, as *V_d_* corresponds to a detectable diameter of 10 mm. Assuming each cancer cell has volume 10^-6^ mm^3^, tumors in our model can grow to a carrying capacity of approximately 12.4 cm in diameter, which is in qualitative agreement with the parameters estimated in [12] (≈ 12.42 cm, assuming a tumor spheroid).

By examining Figures 2(a)-2(b), we clearly observe an improved response to constant therapy when using a phenotype-preserving drug, with a treatment success time *t_c_* nearly seven times as long compared to the resistance-inducing therapy. It can be seen that the tumor composition at treatment conclusion is quite different for each therapy (not shown for this simulation, but see a comparable result in Figures S2(b) and S2(d)), and it appears that the pulsed therapy was not sufficiently strong to hamper the rapid growth of the sensitive population. Indeed, treatment failed quickly due to insufficient treatment intensity in this case, as the population remains almost entirely sensitive. Thus, for this patient under these specific treatments, assuming drug resistance only arises via random stochastic mutations, constant therapy would be preferred. One might argue that a pulsed, equal-magnitude treatment is worse when *α* = 0 simply because less total drug (i.e. AUC) is applied. However, we see that even in this case, intermediate dosages may be optimal (see Figure 4(a) below). Thus, it is not the larger total drug *per se* that is responsible for the superiority of the constant protocol in this case, a point that is reinforced by the fact that the results remain qualitatively unchanged even if the total drug is controlled for (see SI Section C).

**Figure 2:**
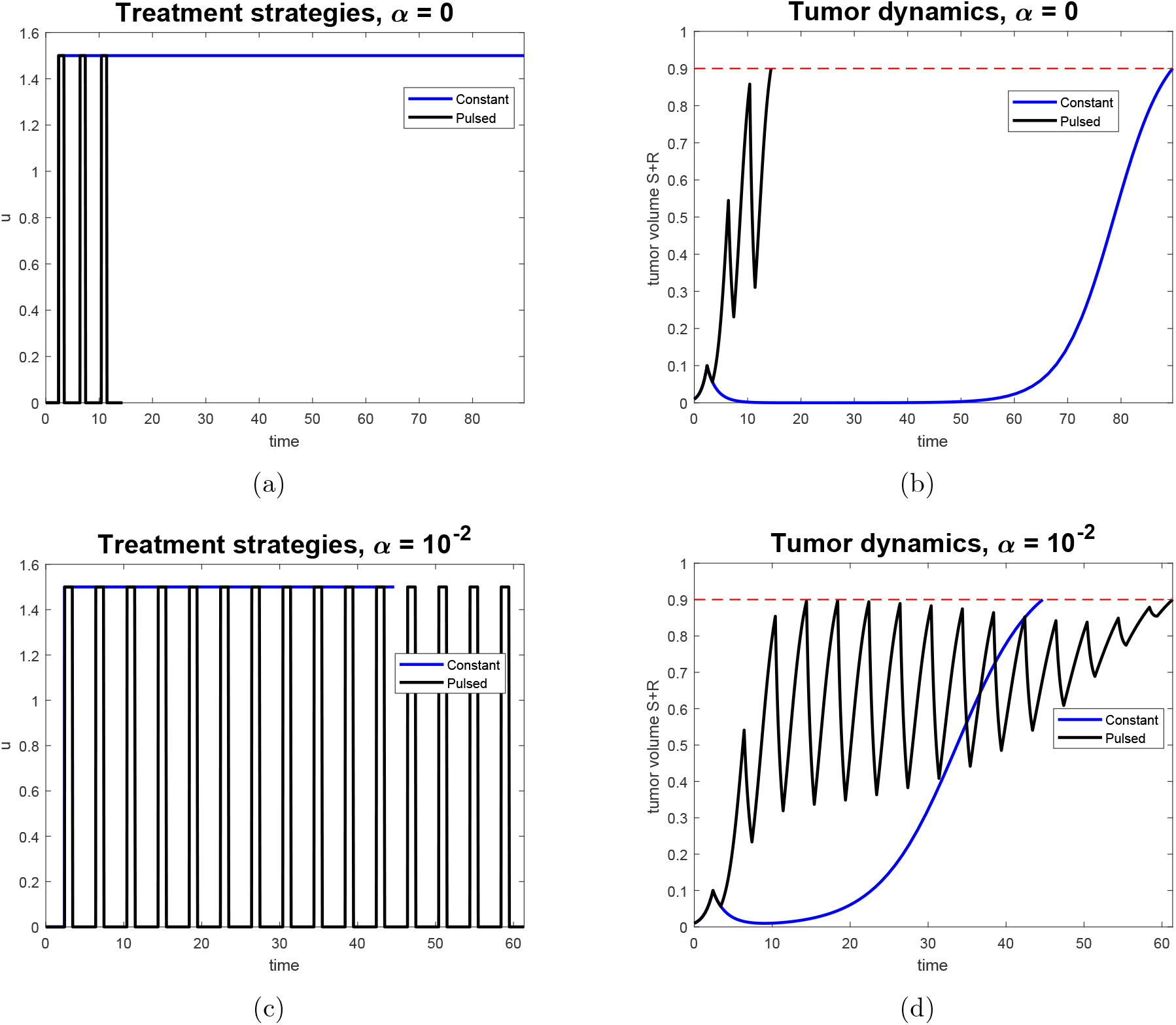
Comparison of treatment efficacy for *phenotype-preserving* (*α* = 0) and resistance-inducing (*α* = 10^-2^) drugs. Left column indicates treatment strategy, while right indicates corresponding tumor volume response. Note that the dashed red line in the right column indicates the tumor volume representing treatment failure, *V_c_*. (a) Constant and pulsed therapies after tumor detection for *α* = 0. (b) Responses corresponding to treatment regimens in (a). (c) Constant and pulsed therapies after tumor detection for *α* = 10^-2^. (d) Responses corresponding to treatment regimens in (c).

Compare this to Figures 2(c)-2(d), which considers the same patient and cytotoxicity, but for a highly inductive drug. The results are strikingly different, and suggest that pulsed therapy is now not worse, and in fact substantially improves patient response (*t_d_* ≈ 61 for pulsed, compared to *t_d_* ≈ 45 for constant). In this case, both tumors are now primarily resistant (see Figures S3(b) and S3(d)), but the pulsed therapy allows prolonged tumor control via sensitive/resistant competitive inhibition. Furthermore, treatment holidays reduce the overall flux into resistance, since the application of the drug itself promotes this evolution. The total amount of drug (AUC) is also smaller for the pulsed therapy (22.5 compared to ≈ 64), so that the pulsed therapy is both more efficient in terms of treatment efficacy, and less toxic to the patient, as side effects are typically correlated with the total administered dose, which is proportional to the AUC.

For these specific parameter values, differences between the constant and pulsed therapy for the inductive drug are not as extensive as in the phenotype-preserving case. However, recall that time has been non-dimensionalized, and hence scale may indeed be clinically relevant. Such differences can be further amplified, and in fact since exact parameters are difficult or even (currently) impossible to measure, qualitative distinctions are paramount. Thus, at this stage, the ranking of therapies, rather than their precise quantitative efficacy, should act as the more important clinical criterion.

From these results, we observe a qualitative difference in the treatment strategy to apply, based entirely on the value of *α*, the degree to which the drug itself induces resistance. Thus, in administering chemotherapy, the resistance-promotion rate *α* of the treatment is a clinically significant parameter. In the next section, we use our model and its output to propose *in vitro* methods for experimentally measuring a drug’s *α* parameter.

### Identifying the Rate of Induced Drug Resistance

The effect of treatment on the evolution of phenotypic resistance may have a significant impact on the efficacy of conventional therapies. Thus, it is essential to understand the value of the induction parameter *α* prior to administering therapy. In this section, we discuss both the theoretical possibility and practical feasibility of determining *α* from different input strategies *u*(*t*).

#### Theoretical Identifiability

We first show that all parameters in system (4)-(5) are identifiable utilizing a relatively small set of controls *u*(*t*) via classical methods from control theory. We provide a self-contained discussion; for a thorough review of theory and methods, see the recent article [72] and the references therein.

Assuming that time and tumor volume are the only clinically observable outputs (i.e. that one cannot readily determine sensitive and resistant proportions in a given population), we measure *V*(*t*) and its derivatives at time *t* = *t_d_* for different controls *u*(*t*). For simplicity, we assume that *t_d_* = 0, so that treatment is initiated with a purely wild-type (sensitive) population. Although the results remain valid if *t_d_* > 0, this assumption will simplify the subsequent computations. For a discussion of the practical feasibility of such methods, see the following section.

Specifically, consider the system (4)-(5) with initial conditions (7). Measuring *V*(*t*) = *S*(*t*) + *R*(*t*) at time *t* = 0 implies that we can identify *S*_0_:

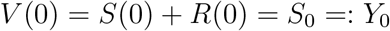

where we adopt the notation *Y_i_*, *i* ≥ 0 for measurable quantities. Similarly, define the following for the given input controls:

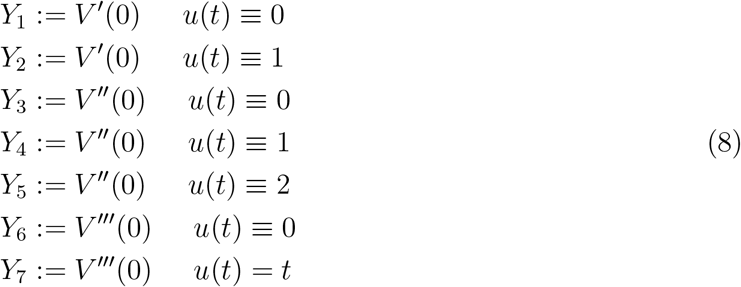

All quantities *Y_i_*, *i* = 0, 1, …, 7 are measurable, as each requires only knowledge of *V*(*t*) in a small positive neighborhood of *t* = 0. Note that the set of controls *u*(*t*) is relatively simple, with *Y*_7_ exclusively determined via a non-constant input.

Each measurable *Y_i_* may also be written in terms of a subset of the parameters *d*, *ϵ*, *p_r_*, and *α*, as all derivatives can be calculated in terms of the right-hand sides of equations (4)-(5). For more details, see SI Section D. Equating the expressions yields a system of equations for the model parameter, which we are able to solve. Carrying out these computations yields the following solution:

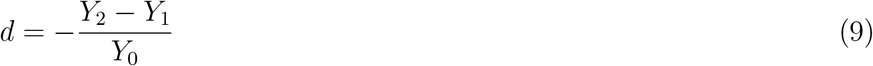

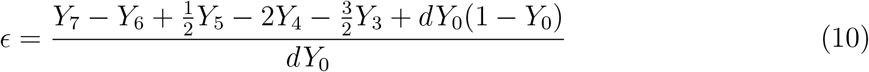

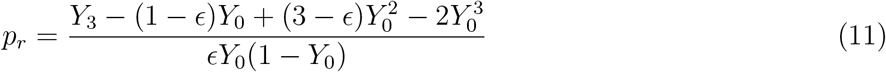

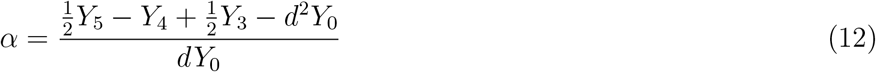

Note that in system (9)-(12), each quantity is determined by the *Y_i_* and the parameter values previously listed; we do not write the solution in explicit form for the sake of clarity, as the resulting equations are unwieldy. Furthermore, the solution of this system relies on the assumption of strictly positive initial conditions (*S*_0_ = *Y*_0_ > 0), wild-type drug induction death rate (*d*), and background mutation rate (*ϵ*), all of which are made in this work.

Equation (12) is the desired result of our analysis. It demonstrates that the drug-induced phenotype switching rate *α* may be determined by a relatively small set of input controls *u*(*t*). As discussed in the previous section, the value of *α* may have a large impact on treatment efficacy, and thus determining its value is clinically significant. Our results now prove that it is possible to compute the induction rate, and hence utilize this information in designing treatment protocols. In the next section, we investigate other qualitative properties that could also be applied to understand the rate of drug-induced resistance.

#### An *in vitro* Experimental Protocol to Distinguish Spontaneous and Drug-Induced Resistance

We have demonstrated that all parameters in (4)-(5) are identifiable, so that it is theoretically possible to determine the phenotype switching rate *α* from a class of input controls *u*(*t*). However, we see that the calculation involved measuring derivatives at the initial detection time *t* = *t_d_*. Furthermore, the utilized applied controls (see equation (8)) are non-constant and thus require fractional dosages to be administered. Clinically, such strategies and measurements may be difficult and/or impractical. In this section, we describe an *in vitro* method for estimating *α* utilizing *constant therapies only*. Specifically, our primary goal is to distinguish drugs with *α* = 0 (phenotype-preserving) and *α* > 0 (resistance inducing). Such experiments, which are described below, may be implemented for a specific drug, even if its precise mechanism of promoting resistance’ remains uncertain.

Before describing the *in vitro* experiment, we note that we are interested in qualitative properties for determining *α*. Indeed, in most modeling scenarios, we have little or no knowledge of precise parameter values, and instead must rely on characteristics that distinguish the switching rate *α* independently of quantitative measurements. Furthermore, as a general framework for drug resistance, the only guaranteed clinically observable output variables are the critical tumor volume *V_c_* and the corresponding time *t_c_* (for a description of the treatment protocol, see above); we cannot expect temporal clonal subpopulation measurements. Assuming *V_c_* is fixed for a given cancer, *t_c_* is thus the only observable that we consider.

By examining (4)-(5), we see that the key parameters dictating the progression to the steady state (*S*, *R*) = (0,1) are d and *α*, as these determine the effectiveness and resistance-induction of the treatment, respectively. Recall that ∊ is the fixed background mutation rate, and p_r_ the relative fitness of the resistant cells. Thus, we perform a standard dose-response experiment for each value of drug sensitivity *d*, and measure the time *t_c_* to reach critical size *V_c_* as a function of *d*. The response *t_c_* will then depend on the applied dosage u (recall that we are only administering constant therapies) and the sensitivity of the wild-type cells *d*, as well as the induction rate *α*:

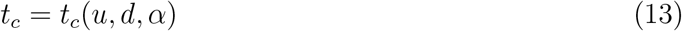

We further imagine that it is possible to adjust the wild-type drug sensitivity *d*. For example, in the case of multi-drug resistance in which the over-abundance of P-glycoprotein affects drug pharmacokinetics, altering the expression of MDR1 via ABCBC1 or even CDX2 [37] may yield a quantifiable relationship between wild-type cell and *d*, thus producing a range of drug-sensitive cell types. Figure 3(a) exhibits a set of dose response curves for representative drug sensitivities *d*, for the case of a resistance-inducing drug (*α* = 10^-2^).

**Figure 3:**
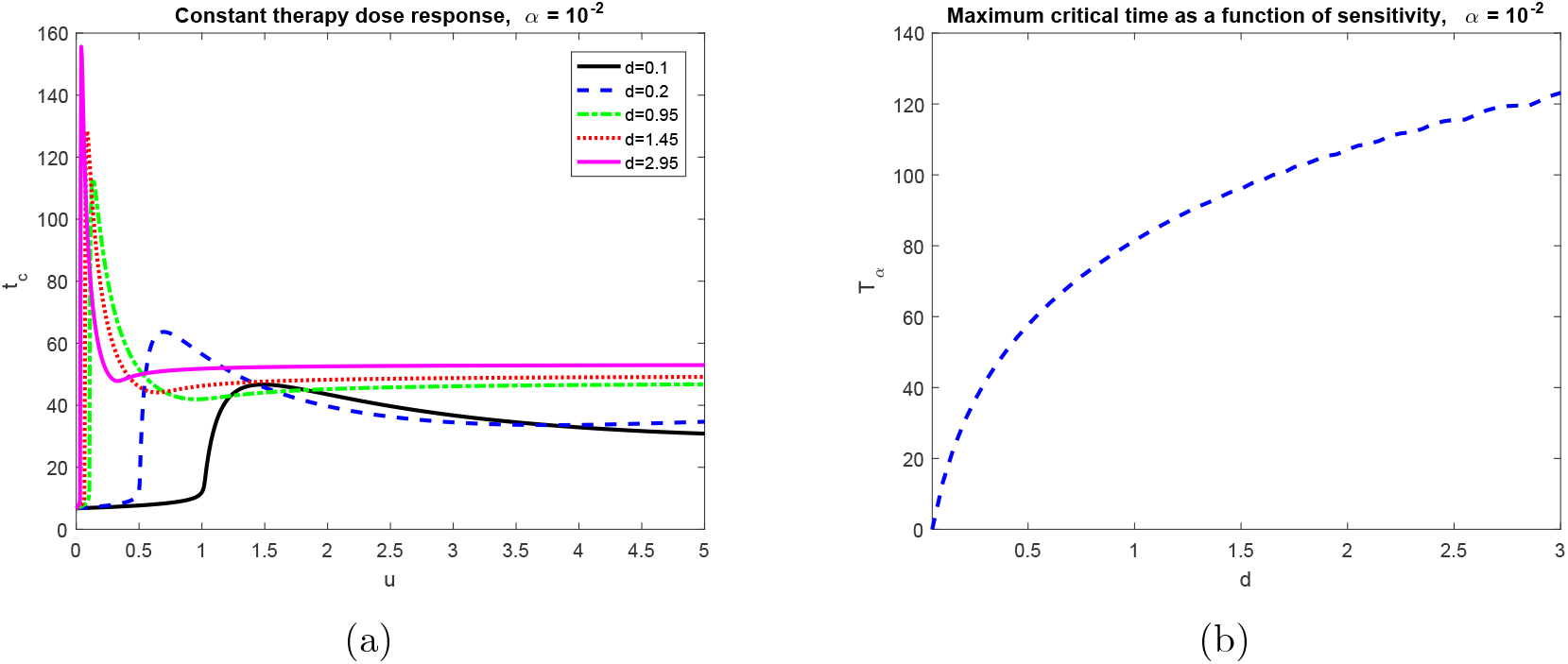
Variation in response time *t_c_* for a treatment inducing resistance. Constant therapy *u*(*t*) ≡ u is applied for *t_d_* ≤ *t* ≤ *t_c_*. Induction rate *α* = 10^-2^, with all other parameters as in Table 1. (a) Time until tumor reaches critical size V_c_ for various drug sensitivities *d*. (*b*) Maximum response time *T_α_*(*d*) for a treatment inducing resistance. Note that time *T_α_*(*d*) increases with drug sensitivity; compare to Figure 4(b) for purely random resistance evolution.

For each of these cell-types, we then define the supremum response time over administered doses:

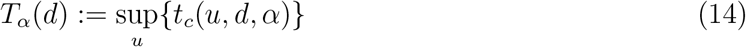

Note that in a laboratory setting, only a finite number of doses will be administered, so that the above supremum is actually a maximum, but for mathematical precision we retain supremum.

Thus, we obtain a curve *T_α_* = *T_α_*(*d*) for each value of the induced resistance rate *α*. We then explore the properties of these curves for different *α* values.

Consider first the case of a phenotype-preserving drug, so that *α* = 0. As *u*(*t*) ≡ *u*, we see that the system (4)-(5) depends only on the product of u and d. Hence, the dependence in (13) becomes of the form *t_c_*(*u* · *d*, 0), and thus the supremum in (14) is instead across the joint parameter *D* := *u* · *d*:

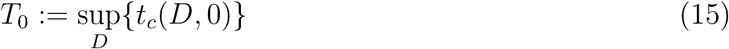

Clearly, this is independent of *d*, so that To is simply a horizontal line for *α* = 0. Qualitatively, the resulting curve will have no variation among the engineered sensitive phenotypes, save for experimental and measurement noise. See Figure 4 for both representative curves (Figure 4(a), comparable to Figure 3(a))) and a plot of *T*_0_ (d) (Figure 4(b)) verifying its independence of *d*. We make two minor technical notes. First, it is important that we assume *d* > 0 here, as otherwise *D* ≡ 0, independent of dose *u* and the supremum is over a one element set. See below for more details and the implications for *α* identifiability. Second, the slight variation for large values of *d* is due to numerical error, as the maximum of *t_c_* occurs at decreasing dosages (see SI Section E and Figure S4 for more details).

**Figure 4:**
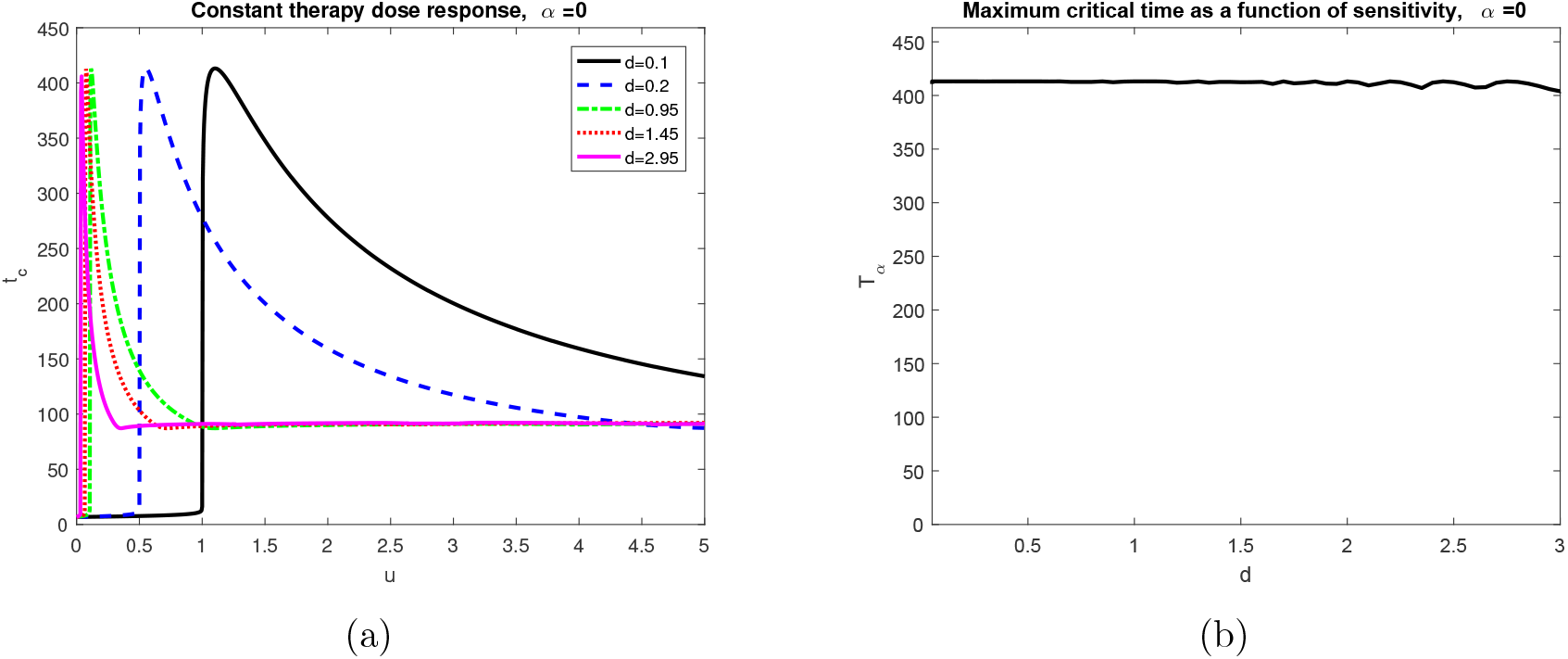
Change in critical time t_c_ for differing drug sensitivities in the case of a phenotype-preserving treatment. (a) Time until tumor reaches critical size *V_c_* for various drug sensitivities *d*. Comparable to Figure 3(a), with *α* = 0. (b) Maximum critical time *T*_0_(*d*). Note that the curve is essentially constant.

Comparing Figures 3(a) and 4(a), we observe similar properties: small *t_c_* for small doses, a sharp increase about a critical *u_c_*, followed by smooth decrease and eventual horizontal asymptote (for mathematical justification, see SI Section E). However, note that for a resistance-inducing drug, (Figure 3(a)), the maximum critical time *T_α_*(*d*) increases as a function of *d*. This is in stark contrast with the constant behavior obtained for *α* = 0, argued above and demonstrated in Figure 4(b). To understand this phenomena further, we plot *T_α_*(*d*) for a fixed induction rate *α* = 10^-2^ in Figure 3(b). The behavior of this curve is a result of the fact that the critical dosage *u_c_* at which *T_α_*(*d*) is obtained is a decreasing function of *d* (see equation (S16) and Figure S4 in SI Section E). But since *u_c_* also controls the amount of resistant cells generated (via the *αu*(*t*)*S* term), resistance growth is impeded by a decreasing *u_c_*. Thus, as a non-negligible amount of resistant cells are necessary to yield *T_α_*(*d*), more time is required for resistant cells to accumulate as *d* increases. Hence, *T_α_*(*d*) increases a function of *d*.

The behavior observed in Figures 3(b) and 4(b) is precisely the qualitative distinction that could assist in determining the induced resistance rate *α*. In the case of a phenotype-preserving drug, the proposed *in vitro* experiment would produce a flat curve, while a resistance-inducing drug (*α* > 0) would yield an increasing function *T_α_*(*d*). Furthermore, we could utilize this phenomena to, in principle, measure the induction rate from the experimental *T_α_*(*d*) curve. For example, Figure 5(a) displays a range of *T_α_*(*d*) for *α* near 10^-2^.

**Figure 5:**
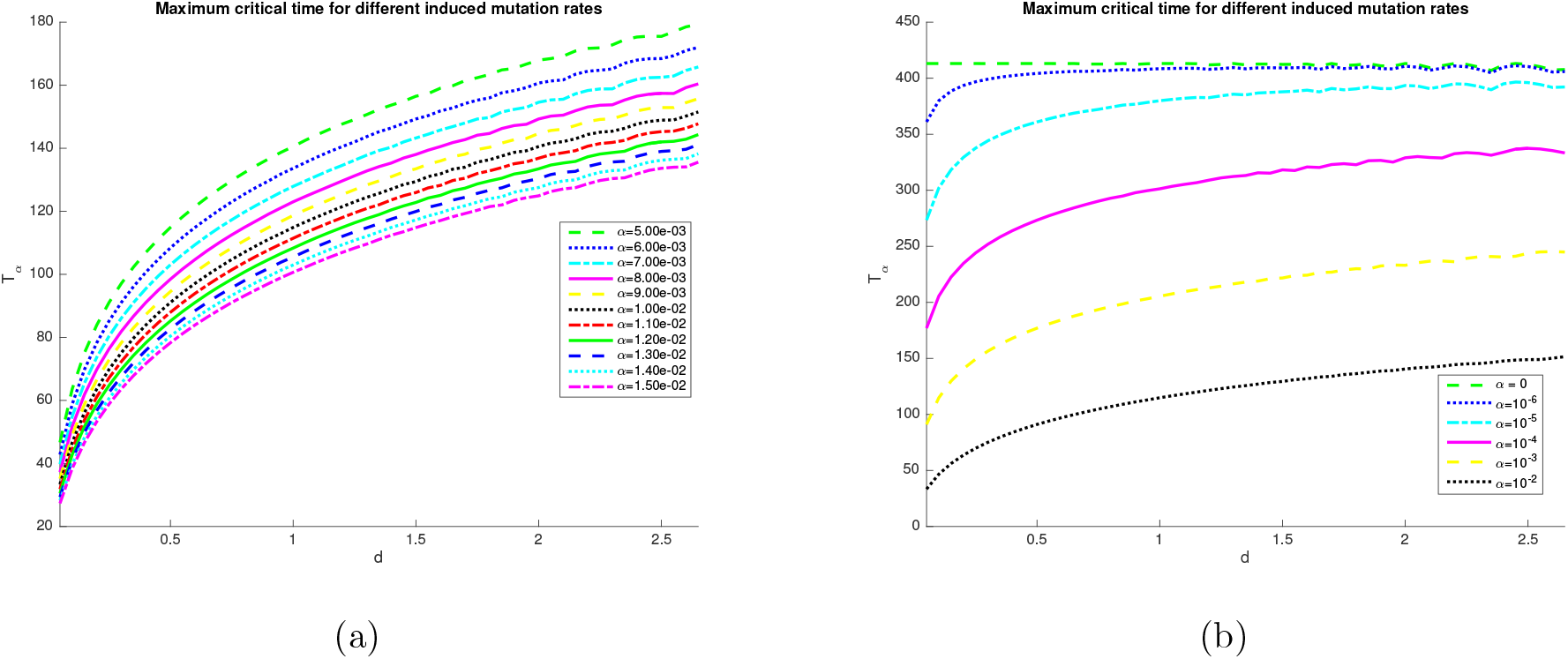
Variation in maximum response time for different induction rates *α*. For details on computation of *T_α_*(*d*), see Figure S4. All other parameters given as in Table 1. (a) Plot of *T_α_*(*d*) for *α* near 10^-2^. (b) Analogous to (a), where *α* is now varied over several orders of magnitude. Non-mutagenic case (*α* = 0) is included for reference.

Figure 5(a) shows a clear dependence of *T_α_*(*d*) on the value of *α*. Quantitatively characterizing such curves would allow us to reverse engineer the induction rate *α*. However, we note that in general the precise characteristics will depend on the other fixed parameter values, such as *p_r_*, *V_c_*, and *ϵ*. Indeed, only order of magnitude estimates may be feasible; illustrative sample curves are provided in Figure 5(b). Two such characteristics are apparent from this figure, both related to the slope of *T_α_*(*d*). First, as *d* → 0^+^, we observe an increase in the slope of *T_α_*(*d*) as *α* decreases (note that in Figure 5(b), only *d* ≥ 0.05 are plotted). This follows from continuity of solutions on parameters and the fact that *T*_0_(*d*) possesses a jump discontinuity at *d* = 0, i.e. its distributional derivative is given by

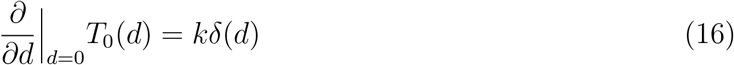

where *δ* is the Dirac function, and k is a positive constant. As discussed previously (see (15) and the subsequent paragraph), *T*_0_(*d*) is flat, except at *d* = 0 where the vector field contains no *u* dependence. Therefore the set over which the maximum is taken is irrelevant, and *T*_0_(*d*) is thus proportional to the Heaviside function, which possesses the distributional derivative (16). The constant *k* is determined by the size of the discontinuity of *T*_0_(*d*). Continuous dependence on parameters then implies that as *α* increases, the resulting derivative decreases away from positive infinity, since the corresponding derivative for *T_α_*(*d*) with *α* > 0 is defined in the classical sense for *α* > 0:

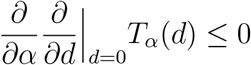

The above argument implies that measuring the slope of *T_α_*(*d*) at *d* = 0 will give a characterization of the phenotypic alteration rate *α* of the treatment. However, such experiments may be impractical, as fine-tuning a cell’s sensitivity near complete resistance may be difficult. Alternatively, one could analyze the degree of flatness for a relatively large *d* (so to be sufficiently far from *d* = 0) and correlate this measure with *α*. For example, examining *d* = 2 in Figure 5(b), we see that the relative slope of *T_α_*(*d*) with respect to *d* should correlate with decreasing *α*. A similar argument to the above makes this rigorous, for *d* sufficiently large. Practical issues still arise, but this second method does provide a more global method for possibly computing *α*. Indeed, slopes at a given *d* can be approximated by a wider range of secant approximations, as the result holds for a range of *d*, as compared to the previously-discussed case when *d* is near zero. Furthermore, our focus is largely on the qualitative aspects of *α* determination (such as the differences in Figures 3(a) and 4(a)) and determining whether the treatment itself induces resistance to emerge.

### Conclusions and Future Work

In this work, we have analyzed two distinct mechanisms that can result in drug resistance. Specifically, a mathematical model is proposed which describes both the spontaneous generation of resistance, as well as drug-induced resistance. Utilizing this model, we contrasted the effect of standard therapy protocols, and demonstrated that contrary to the work in [48], the rate of resistance induction may have a significant effect on treatment outcome. Thus, understanding the dynamics of resistance evolution in regards to the applied therapy is crucial.

To demonstrate that one can theoretically determine the induction rate, we performed an identifiability analysis on the parameter *α*, and showed that it can be obtained via a set of appropriate perturbation experiments on *u*(*t*). Furthermore, we presented an alternative method, utilizing only constant therapies, for understanding qualitative differences between the purely spontaneous and induced cases. Such properties could possibly be used to design *in vitro* experiments on different pharmaceuticals, allowing one to determine the induction rate of drug resistance without an *a priori* understanding of the precise mechanism. We do note, however, that such experiments may still be difficult to perform in a laboratory environment, as engineering cells with various drug sensitivities *d* may be challenging. Indeed, this work can be considered as a thought experiment to identify qualitative properties that the induction rate *α* yields in our modeling framework.

Our simple model allows significant insight into the role of random versus induced resistance. Of course, more elaborate models can be studied by incorporating more biological detail. For example, while our two-equation model classifies cells as either sensitive or resistant, not all resistance is treated equally. Some resistant cancer cells are permanently resistant, whereas others could transition back to a sensitive state [65]. This distinction may prove to be vitally important in treatment design. A possible extension of our model is one in which we distinguish between sensitive cells *S*, non-reversible resistant cells *R_n_*, and reversible resistant cells *R_r_*:

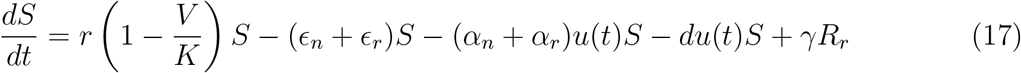

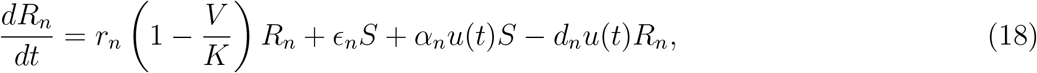

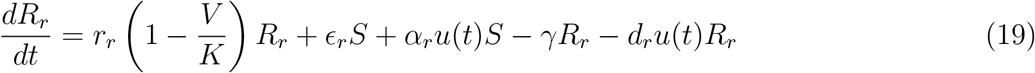

Here *V* denotes the entire tumor population, i.e.

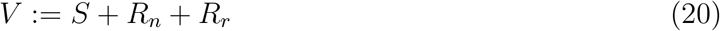

In this version of the model, the non-reversible resistant cells *R_n_* can be thought of as resistant cells that form via genetic mutations. Under this assumption, *ϵ_n_* represents the rate at which spontaneous genetic mutations give rise to resistance, and *α_n_* is the drug-induced resistance rate. This situation can be classified as non-reversible since it is incredibly unlikely that genomic changes that occur in response to treatment would be reversed by an “undoing” mutation. Therefore, once cells confer a resistant phenotype through an underlying genetic change, we assume they maintain that phenotype. This term could also be thought of as describing resistance that forms via stable epigenetic alterations, or resistance that forms via some combination of genetic and stable epigenetic changes.

On the other hand, reversible resistant cells *R_r_* denote resistant cells that form via phenotype-switching, as described in [65]. Random phenotype-switching in the absence of treatment is captured in the *ϵ_r_S* term. This is consistent (and indeed necessary) to understand the experimental results in [65], where a stable distribution of MDR1 expressions is observed even in the absence of treatment. The *α_r_u*(*t*)*S* term represents the induction of a drug-resistant phenotype. Phenotype-switching is often reversible, and therefore we allow a back transition from the Rr compartment to the sensitive compartment at a non-negligible rate *γ* [65] (see SI Section B. Formulated in this way, the model can be calibrated to experimental data and we can further consider the effects of the dosing strategy on treatment response. We plan to further study this model in future work.

Overcoming drug resistance is crucial for the success of both chemotherapy and targeted therapy. Furthermore, the added complexity of induced drug resistance complicates therapy design, as the simultaneous effects of tumor reduction and resistance propagation confound one another. Mathematically, we have presented a clear framework for differentiating random and drug-induced resistance, which will allow for clinically actionable analysis on a biologically subtle, yet important, issue.

## Supporting Information (SI)

The supplement contains additional results and extensions related to the model (1)-(2) presented in the main text. Sections include details on the mathematical characteristics of solutions of system (4)-(5), an extension describing the reversibility of drug resistance, treatment regimes with normalized dosages, details on structural identifiability, and an expanded discussion on the maximum critical time *T_α_*(*d*) (see (14)).

### A Fundamental Solution Properties of Resistance Model

For convenience, system (4)-(5) is reproduced below:

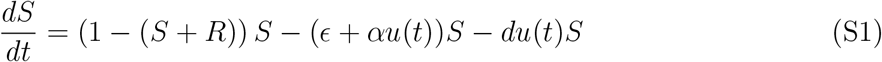

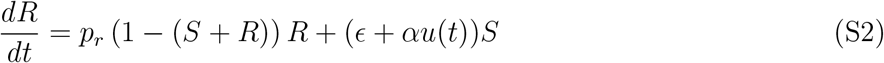

We begin with a standard existence/uniqueness result, as well as the dynamical invariance of the triangular region

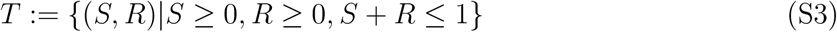

Note that *T* represents the region of non-negative tumor sizes below 1. Biologically, this implies that all solutions remain physical (non-negative) and bounded above by the carry capacity (non-dimensionalized to 1 here, generally *K* in (1)-(2)).

Theorem 1. *For any bounded measurable control u* : [0, ∞) → [0, *u*_max_], *with u*_max_ < ∞, *and* (*S*_0_, *R*_0_) ∈ *T*, *the initial value problem* (S1)-(S2),

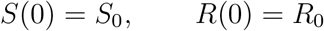

*has a unique solution* (*S*(*t*), *R*(*t*)) *defined for all times* 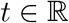. *Furthermore, under the prescribed dynamics, region T is invariant*.

#### Proof

Existence and uniqueness of local solutions follow from standard results in the theory of differential equations; see for example Theorem 2.1.1 and 2.1.3 in [5]. Since the vector field

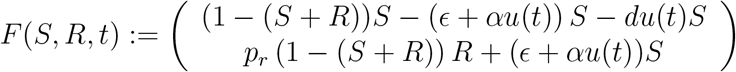

is analytic for 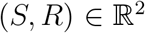, the existence of maximal solutions defined for all 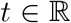 will follow from boundedness (see Theorem 2.1.4 in [5]), which we demonstrate below.

Uniqueness implies that solutions remain in the first quadrant for all *t* ≥ 0. Indeed, we first note that (0, 0) and (0, 1) are steady states for any control *u*(*t*). As *S* =0 implies that *Ṡ* = 0, we see that the *R*-axis in invariant, with 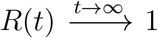. Similarly, *R* =0 implies *Ṙ* ≥ 0, and hence all trajectories with (*S*_0_, *R*_0_) ∈ *T* remain non-negative for all *t*. As *V* = *S* + *R* satisfies the differential equation

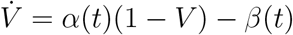

where *α*(*t*) := *S*(*t*) + *p_r_R*(*t*), *β*(*t*) := *du*(*t*)*S*(*t*) are both non-negative, *V*_0_ < 1 ⇒ *V*(*t*) < 1. Thus, if the initial conditions (*S*_0_, *R*_0_) reside in *T*, we are guaranteed that (*S*(*t*), *R*(*t*)) ∈ *T* for all time *t*, as desired.

We now prove that asymptotically the cells will evolve to become entirely resistant. For simplicity, we assume that the tumor is initially below carrying capacity, although a similar result holds for *V*_0_ > 1.

Theorem 2. *For any bounded measurable control u* : [0, ∞) → [0, *u*_max_], *with u*_max_ < ∞, *and initial conditions* (*S*_0_, *R*_0_) ∈ *T*, *solutions of system* (S1)-(S2) *will approach the steady state*(*S*, *R*) = (0, 1):

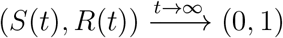

#### Proof

From Theorem 1, we have that 0 ≤ *S*(*t*) + *R*(*t*) ≤ 1, so that (S2) implies

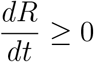

As 0 ≤ *R*(*t*) ≤ *S*(*t*) + *R*(*t*) ≤ 1, *R*(*t*) must converge, so that there exists 0 ≤ *R*_*_ ≤ 1 such that

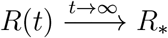

By uniqueness, for all admissible controls *u*(*t*), the line *R* = *R*_*_ is attracting and invariant, and thus the corresponding sensitive component *S*(*t*) along this line must satisfy

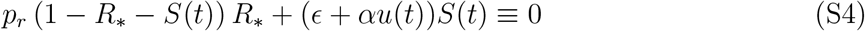

Since *ϵ* > 0 and *u*(*t*) ≥ 0, and both terms on the left-hand side of (S4) are non-negative, this is only possible if *S*(*t*) ≡ 0, *R*_*_ = 1. Hence the line *R* = *R*_*_ is in actuality the point (*S*, *R*) = (0, 1), so that 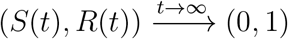 for all (*S*_0_, *R*_0_) ∈ *T*, as desired.

### B Reversible phenotype switching

In the model analyzed in this work (system (4)-(5)), we assumed that resistance is non-reversible. However, experiments suggest [64] that phenotypic alterations are generally unstable, and hence a non-negligible back transition exists. In this appendix, we demonstrate that an extension of our model to include this phenomenon does not change the qualitative results presented previously, at least for parameter values consistent with experimental data.

To model reversible drug resistance, we include a constant per-capita transition rate from the resistant compartment *R* back to the wild cell-type *S*. Denoting this rate by *γ*, we obtain the system

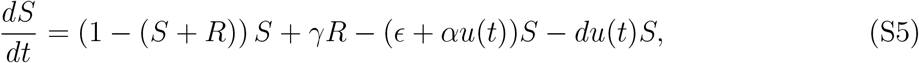

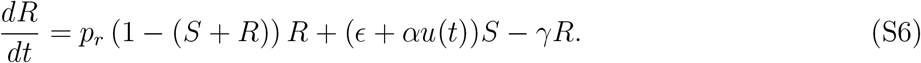

We first consider an appropriate value of the rate *γ*. Note that in the absence of treatment (*u*(*t*) ≡ 0), the system becomes

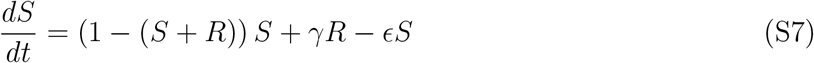

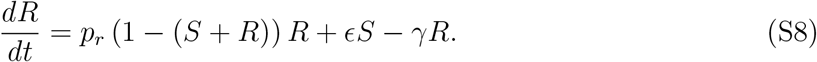

Note that the tumor volume *V*(*t*) = *S*(*t*) + *R*(*t*) satisfies the equation

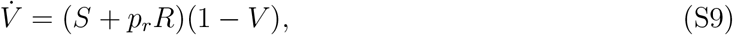

which is non-decreasing (recall Theorem 1). This implies that the system approaches a steady-state (*S*_*_, *R*_*_), which can be easily computed as

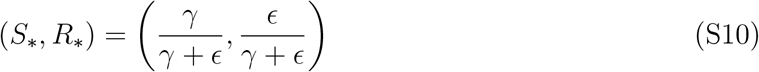

Note that (S10) lies on the line *V* = 1.

Pisco and colleagues [64] measure a 1 – 2% subpopulation of clonally derived HL60 cells which consistently express high levels of MDR1, which we equate with the resistant population *R*. Using the 2% upper bound, this then implies that

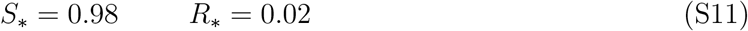

Solving equations (S10) with the above values then determines the ratio 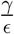:

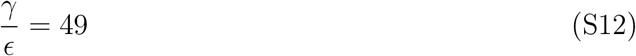

We now repeat the constant vs. pulsed experiments discussed in the main text, but for the reversible system (S5)-(S6). Parameter values are taken again as in Table 1, and *γ* is determined via (S12). Results are presented in Figure S1, and should be compared to Figure 2. Note that the same qualitative (and indeed quantitative) conclusions hold: constant therapy improves response time compared to pulsing when *α* = 0, while the reverse is true for *α* = 10^-2^. Thus, including instability of the resistant cell subpopulation still implies that knowledge of the resistance-induction rate *α* for a chemotherapy is critical when designing therapies. We note that precise agreement of Figures S1 and 2 is due to the small values taken for ∊ and hence *γ*.

**Figure S1:**
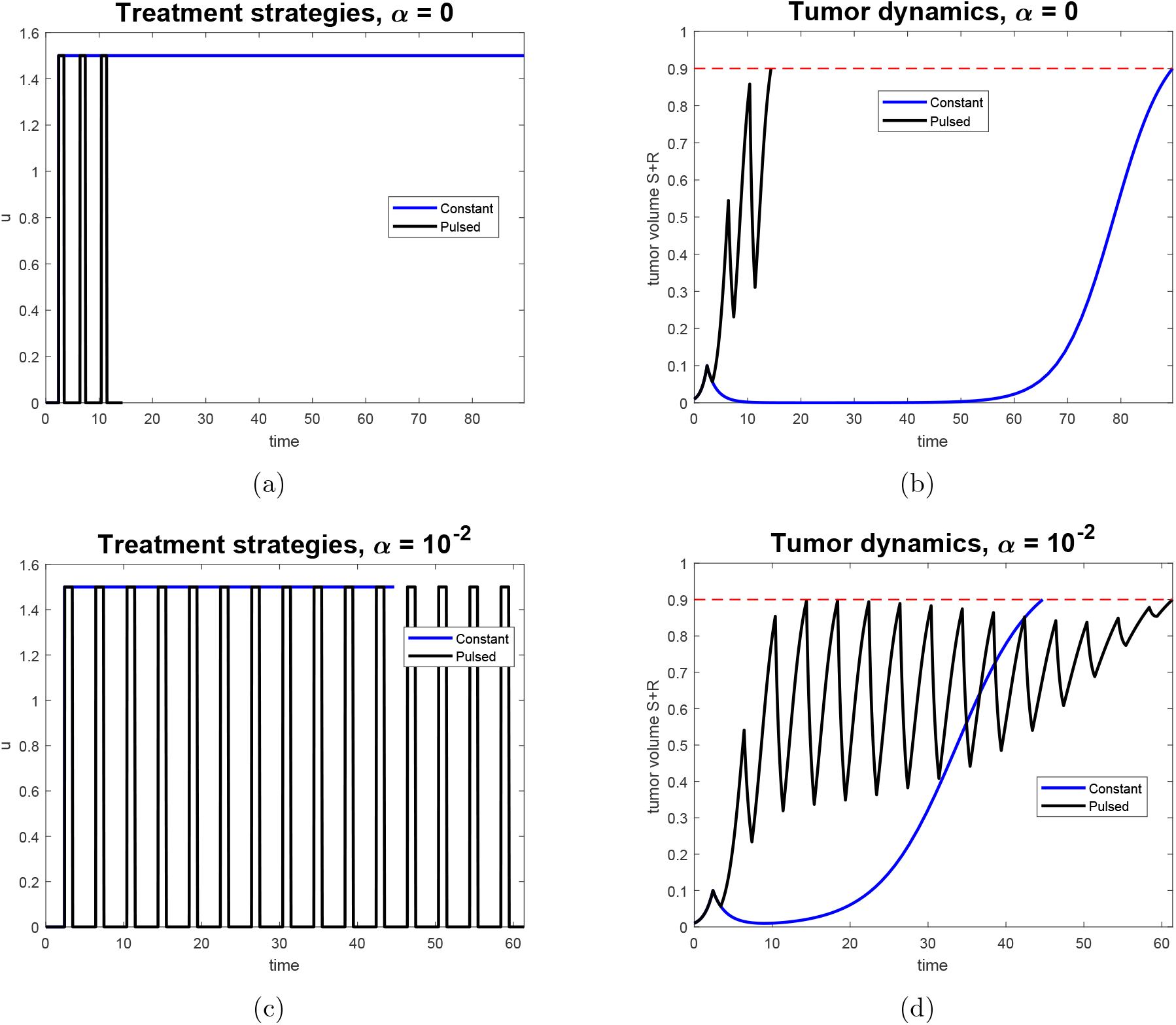
Comparison of treatment efficacy for *phenotype-preserving* (*α*= 0) and resistance-inducing (*α* = 10^-2^) drugs, where *resistance is reversible*. Left column indicates treatment strategy, while right indicates corresponding tumor volume response. Note that the dashed red line in the right column indicates the tumor volume representing treatment failure, *V_c_*. (a) Constant and pulsed therapies after tumor detection for *α* = 0. (b) Responses corresponding to treatment regimens in (a). (c) Constant and pulsed therapies after tumor detection for *α* = 10^-2^. (d) Responses corresponding to treatment regimens in (c).

### C Treatment comparison for equal AUC

Here we provide an analogous comparison of treatment outcomes between constant and pulsed therapy as in the main text. However, treatment magnitudes *u*_on,*c*_ and *u*_on,*p*_ are chosen such that

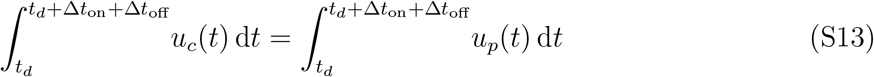

which is equivalent to the conservation of total administered dose between both strategies over a single pulsing cycle. In (S13), *u_c_*(*t*) and *u_p_*(*t*) denote the constant and pulsed therapy schedules, respectively. The constant therapy magnitude *u*_on,*c*_ is fixed (arbitrarily) at 0.5, which by (S13) implies that *u*_on,*p*_ = 5. We also adjust Δ*t*_on_ = 0.5, Δ*t*_off_ = 4.5, and all other parameter values remain as in Table 1.

**Figure S2:**
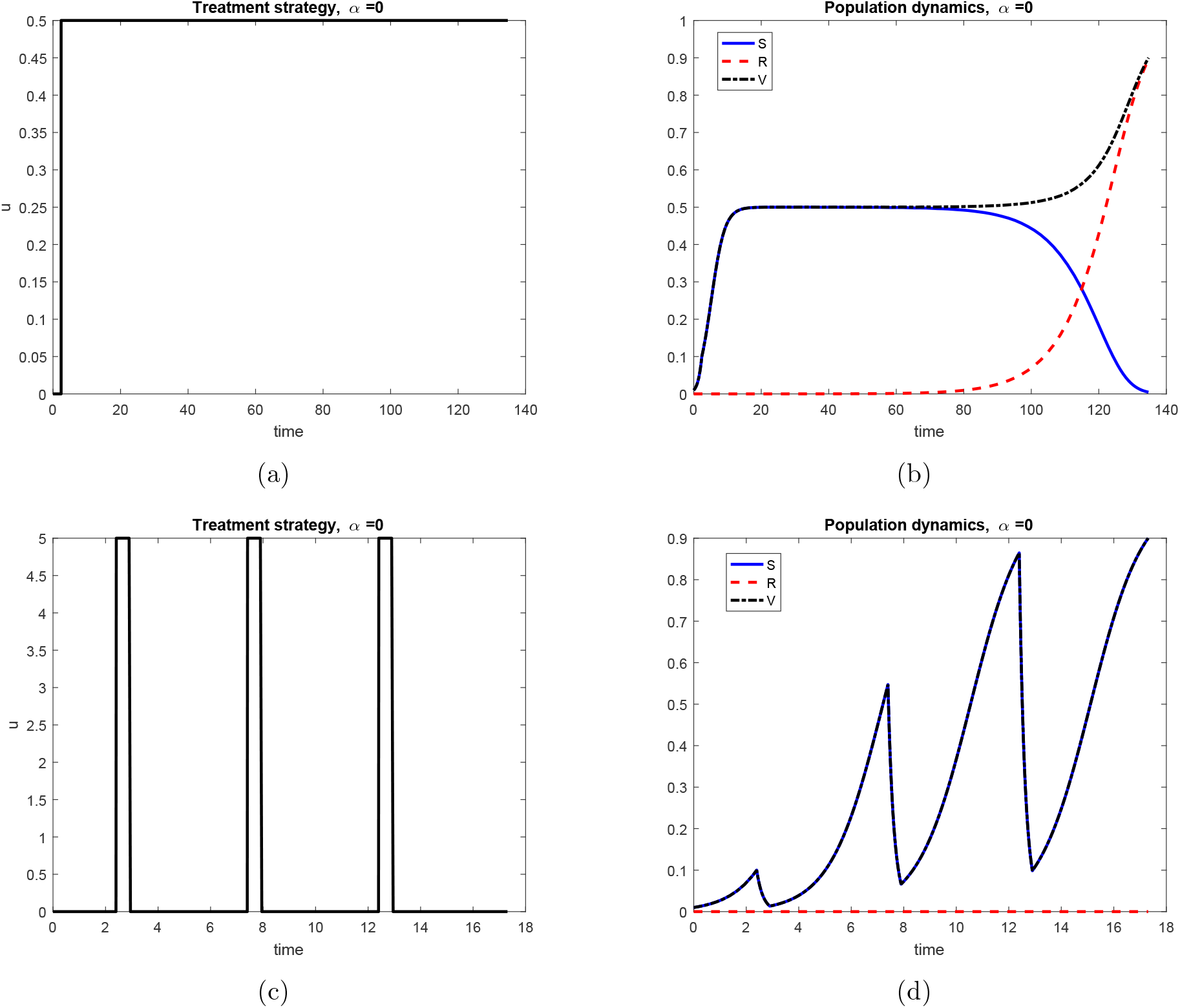
Treatment dynamics for phenotype-preserving drug (*α* = 0). Left column indicates treatment strategy, while right indicates corresponding population response. (a) Constant treatment after tumor detection. (b) Response to constant treatment. Note that at the time of treatment failure, the tumor is essentially entirely resistant. (c) Pulsed therapy after tumor detection. (d) Response to pulsed therapy. Note that the treatment fails much earlier than for the constant dosage, and that the tumor is primarily drug sensitive.

Results of the simulations are presented in Figures S2 and S2. Note that here Figure S2 displays the results for the phenotype-preserving drug (*α* = 0), while Figure S3 represents the resistance-inducing drug (*α* = 10^-2^). Each drug is simulated for the two distinct strategies; each strategy is represented in the left column of the respective figures. The right column illustrates the population response, for both the individual populations (sensitive *S* and resistant *R* cells), as well as for the total tumor volume *V* = *S* + *R*. As previously discussed, treatment is continued until a critical tumor size *V_c_* is obtained, and the corresponding time *t_c_* is utilized as measure of treatment efficacy, with a larger *t_c_* indicating a better response.

**Figure S3:**
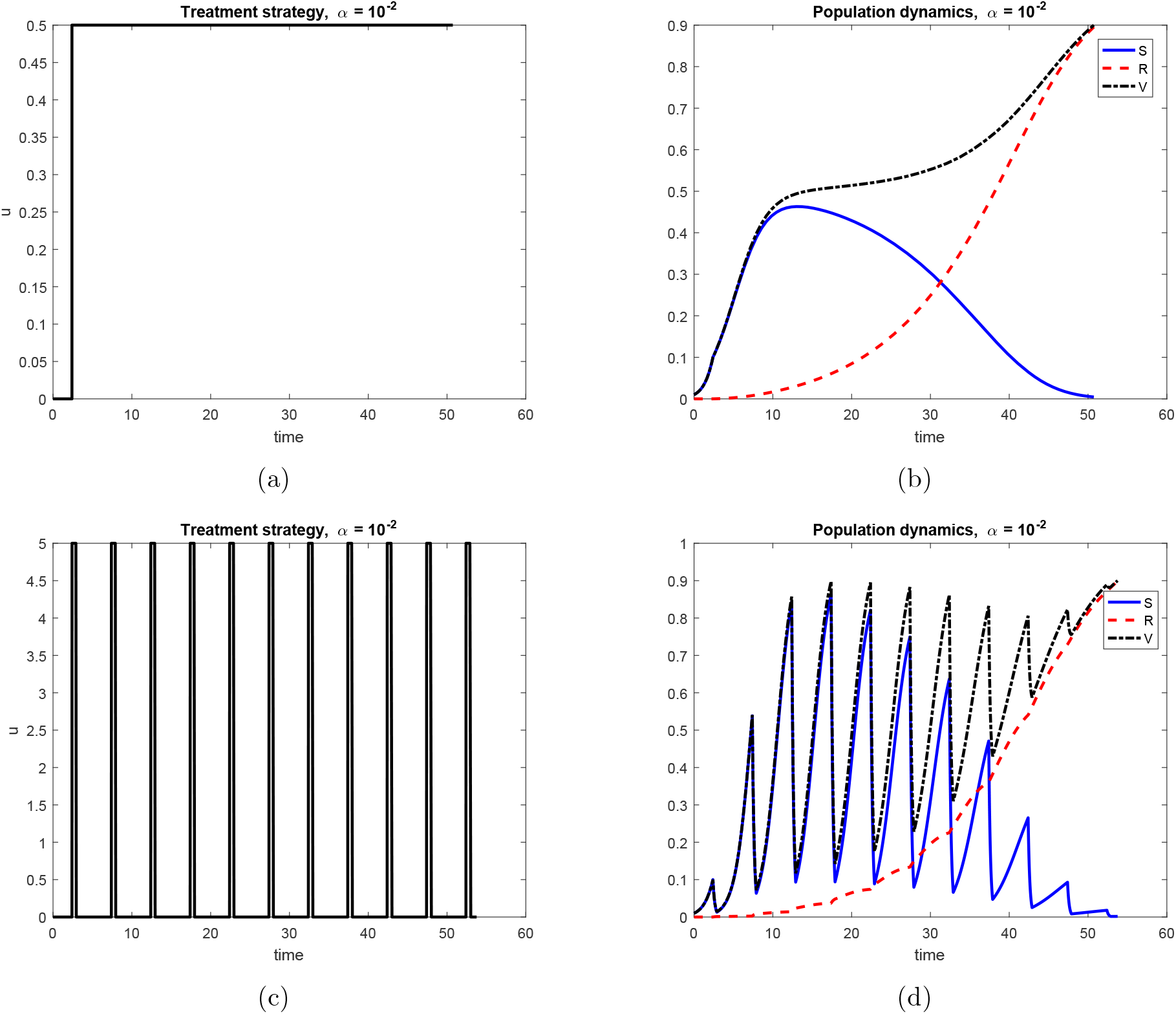
Treatment dynamics for resistance-inducing drug (*α*= 10^-2^). Left column indicates treatment strategy, while right indicates corresponding population response. (a) Constant treatment after tumor detection. (b) Response to constant treatment. Note that at the time of treatment failure, the tumor is essentially entirely resistant. Similar dynamics to Figure 2(b), although with a shorter survival time. (c) Pulsed therapy after tumor detection. (d) Response to pulsed therapy. Note that here, in contrast to case of a phenotype-preserving drug as shown in Figure S2, pulsed therapy exhibits a longer survival time.

Our results are qualitatively in agreement with those presented in the main text (Figure 2), where no preservation of total administered drug (AUC) was considered. Specifically, we observe a superior response with constant therapy for the phenotype-preserving drug (*α* = 0), while the situation is reversed for the resistance inducing drug (*α* = 10^-2^).

### D Identifiability analysis

In this section, we provide additional details on the theoretical identifiability of model parameters. As mentioned in the main text, all higher-order derivatives at initial time *t* = 0 may be calculated in terms of the initial conditions (*S*(0), *R*(0)) and the control function *u*(*t*). For example, for an arbitrary system

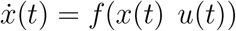

with external control *u*(*t*), the second derivative *ẍ* may calculated using the chain rule:

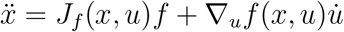

where *J_f_*(*x*) is the Jacobian matrix of *f*, evaluated at state *x* and control *u*. If *x*(0) = *x*_0_ is known, the above expression is a relation among parameters, together with *u* and 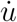 evaluated at time *t* = 0. An analogous statement holds for a measurable output *y* = *h*(*x*), but will involve the Jacobian of *h* as well. Concretely, for the model of induced drug resistance (4)-(5), first derivatives of the tumor volume may be calculated as

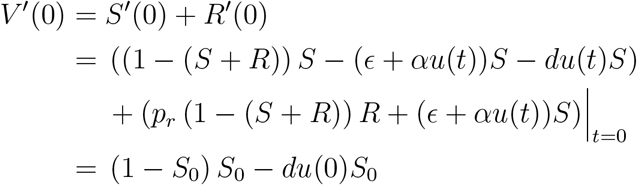

for any control *u*(*t*) (recall that *R*(0) = 0). Similarly, for the second derivative, we compute:

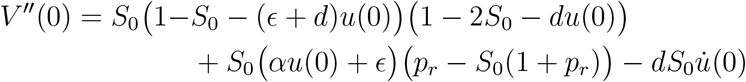

Using such expressions (or, more precisely, the Lie derivatives of the vector field, see [72]) for the controls in (8), one is able to obtain a set of equations between the set of *Y_i_*, *i* =0, 1, …, 7 and the parameters *d*, *ϵ*, *p_r_*, and *α*. Solving these equations allows us to determine the parameters with respect to the measurable quantities. The algebraic solution is system (9)-(12).

### E Analysis of critical time *T_α_*(*d*)

We provide a qualitative understanding of the properties of *T_α_*(*d*), the maximum time, across all constant dosages, for the tumor to reach size *V_i_*. This appendix is designed to explain the basic properties discussed in the *In Vitro* Identifiability section of the main text.

We first note that *T_α_*(*d*) is achieved at a medium dosage *u_c_*. More precisely, we describe the qualitative properties of Figures 3(a) and 4(a). Fix a drug sensitivity *d*. For small *u*, the sensitive subpopulation is not sufficiently inhibited, and hence expands rapidly to cross the threshold *V_c_*, with an essentially homogenous population of sensitive cells. Indeed, as *u* → 0, the dynamics of (S1)-(S2) approach those of

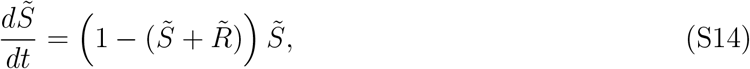

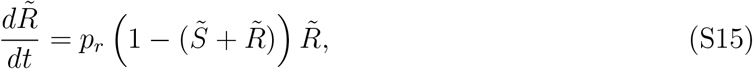

for small finite times, as *ϵ* ≪ 1. Trajectories of (S14)-(S15) remain on the curve

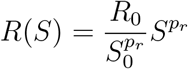

as the solution approaches the line *S* + *R* = 1. The critical time *t_c_* is then determined by the intersection of this curve with *S* + *R* = *V_c_*, and hence has sensitive population *S_c_* at *t_c_* given by the unique solution in the first quadrant of

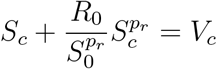

As *R*_0_ ≪ 1 (since *ϵ* ≪ 1), *S_c_* ≈ *V_c_*, as claimed. Time *t_c_* is thus small for small u.

Increasing values of *u* imply *t_c_* to also increase, since the overall growth rate of the sensitive cells is decreased. However, there exists a critical dose *u_c_* such that sensitive cells alone are not able to multiply sufficiently to attain Vc, so that the critical volume must have non-negligible contributions from the resistant fraction. This then leads to the bifurcation apparent in Figures 3(a) and 4(a). We can even approximate the critical dosage maximizing *t_c_*, as *V_c_* must be an approximation for the carrying capacity of the sensitive cells:

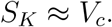

By examining the right-hand side of (S1), and assuming that the dynamics of the resistant population are negligible (which is accurate in the early stages of treatment; see Figures 2(b) and 3(b)) we see that the dose yielding the maximum temporal response should be

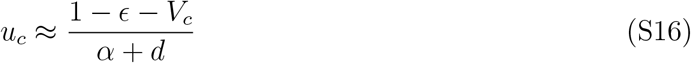

That is, the dosage where *T_α_*(*d*) is obtained is given approximately by the expression in (S16). For a sample numerical comparison of the predicted formula (S16) and a numerical optimization over a range of drug sensitivities *d*, see Figure S4. Note that in actuality, *S_K_* < *V_c_*, as the resistant dynamics cannot be ignored entirely. Thus, the precise value of *u_c_* will be smaller than that provided in the previous formula, as we numerically observe. Lastly, *u_c_* decreases with increasing values of parameter *d*, and thus requires an increasingly fine discretization to numerically locate the maximum value. Hence, some numerical error is observed in Figure 4(b).

**Figure S4:**
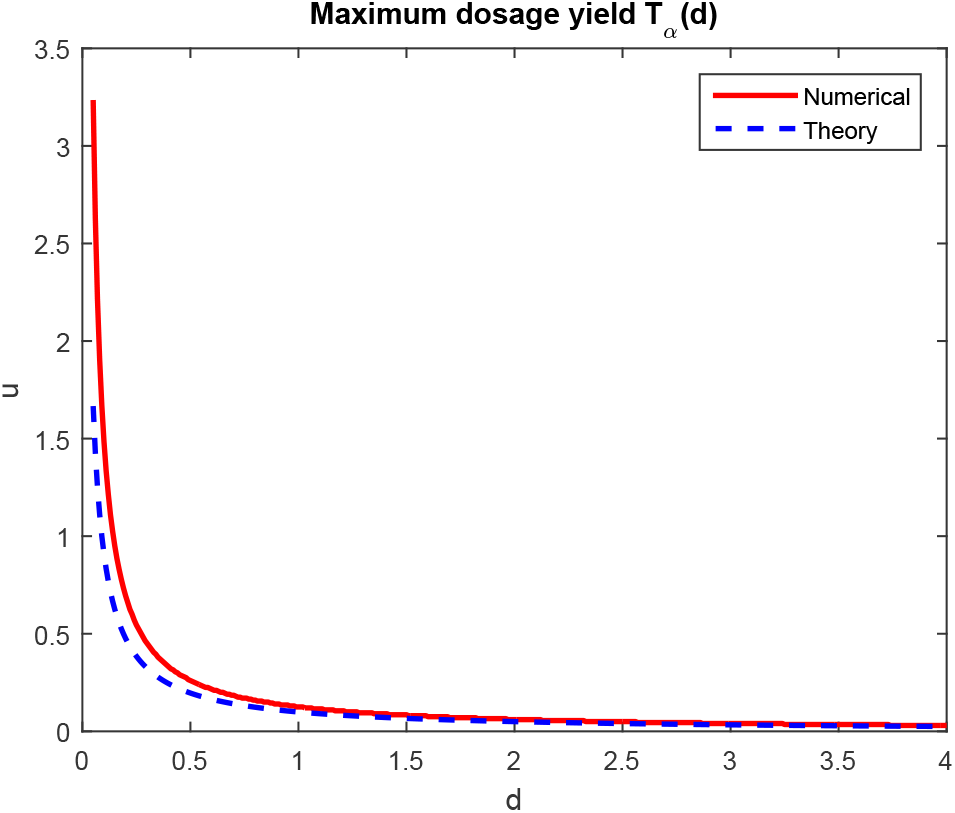
Dose yielding maximal response time *T_α_*(*d*) computed numerically, as well as the approximation given by (S16). All parameters appear as in Table 1, and *α* = 10^-2^. The numerical maximum is computed over a discretization of constant dosage procedures *u*(*t*) ≡ *u*, for *u* ∈ [0, 5], with a mesh size Δ*u* = 0.005.

Lastly, as *u* is increased further, the dosage becomes sufficiently large so that the inhibition of *S* via therapy implies that S cannot approach the critical volume *V_c_*, and hence *V_c_* is again reached by an essentially homogeneous population, but of resistant cells. Since resistant cells divide at a slower rate (*p_r_* < 1), the corresponding time *t_c_* is smaller. For a schematic of the three regimes described above, see Figure S5.

**Figure S5:**
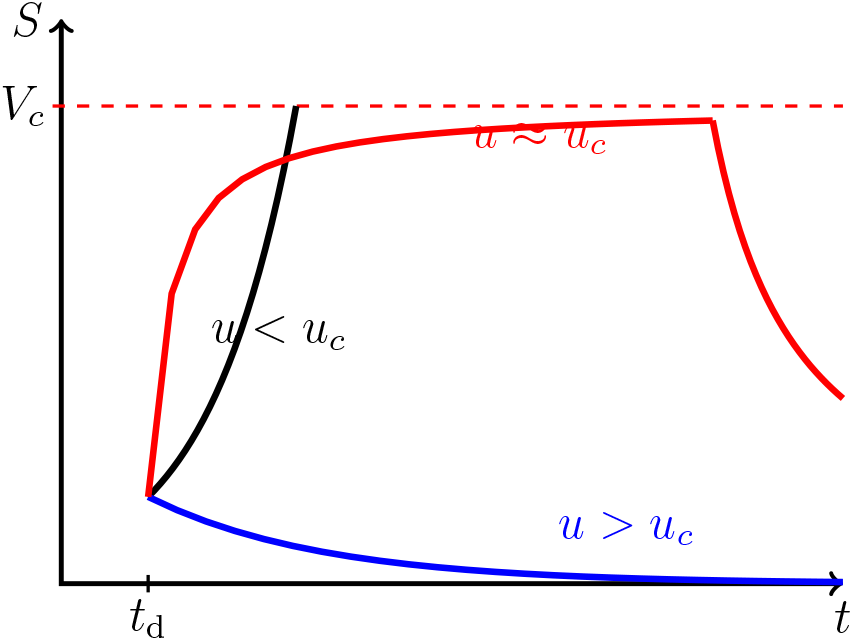
Schematic demonstrating dynamics of variation in *t_c_* on dosage *u*. Sensitive cell population plotted as a function of time, for three representative doses. For *u* < *u_c_*, sensitive cells grow and reach *V_c_* in a short amount of time. As 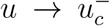, the sensitive population approaches its approximate carrying capacity of *V_c_*, but subsequently decreases due to the dynamics of resistance. Here *t_c_* is maximized, as the sensitive population spends a large amount of time near *V_c_*. For *u* > *u_c_*, the sensitive population is eliminated quickly, and Vc is obtained by a primarily resistant population.

## References

[1] D. Axelrod, S. Vedula, and J. Obaniyi. Effective chemotherapy of heterogeneous and drug-resistant early colon cancers by intermittent dose schedules: a computer simulation study. Cancer Chemother. Pharmocol., 79:889–898, 2017.

[2] B. C. Bender, E. Schindler, and L. E. Friberg. Population pharmacokinetic–pharmacodynamic modelling in oncology: a tool for predicting clinical response. British journal of clinical pharmacology, 79(1):56–71, 2015.

[3] I. Bozic and M. Nowak. Timing and heterogeneity of mutations associated with drug resistance in metastatic cancers. Proceedings of the National Academy of Sciences, 111:15964–15968, 2014.

[4] I. Bozic, J. Reiter, B. Allen, T. Antal, K. Chatterjee, P. Shah, Y. S. Moon, A. Yaqubie, N. Kelly, D. Le, E. Lipson, P. Chapman, L. Diaz, B. Vogelstein, and M. Nowak. Evolutionary dynamics of cancer in response to targeted combination therapy. eLife, 2:1–15, 2013.

[5] A. Bressan and B. Piccoli. Introduction to mathematical control theory. AIMS series on applied mathematics, Philadelphia, 2007.

[6] K. R. Brimacombe, M. D. Hall, D. S. Auld, J. Inglese, C. P. Austin, M. M. Gottesman, and K.-L. Fung. A dual-fluorescence high-throughput cell line system for probing multidrug resistance. Assay and drug development technologies, 7(3):233–249, 2009.

[7] T. Brocato, P. Dogra, E. J. Koay, A. Day, Y.-L. Chuang, Z. Wang, and V. Cristini. Understanding drug resistance in breast cancer with mathematical oncology. Curr. Breast Cancer Rep., 6:110–120, 2014.

[8] M. Carlino, C. Fung, H. Shahheydari, J. Todd, S. Boyd, M. Irvine, A. Nagrial, R. Scolyer, R. Kefford, G. Long, and H. Rizos. Preexisting MEK1P124 mutations diminish response to BRAF inhibitors in metastatic melanoma patients. Clinical Cancer Research, 21:98–105, 2015.

[9] C. Carrère. Optimization of an in vitro chemotherapy to avoid resistant tumors. Journal of Theoretical Biology, 413:24–33, 2017.

[10] S. Chakrabarti and F. Michor. Pharmacokinetics and drug-interactions determine optimum combination strategies in computational models of cancer evolution. Cancer Research, To appear, 2017.

[11] M. Chapman, T. Risom, A. Aswani, R. Dobbe, R. Sears, and C. Tomlin. A model of phenotypic state dynamics initiates a promising approach to control heterogeneous malignant cell populations. In 2016 IEEE 55th Conference on Decision and Control (CDC), pages 2481–2487. IEEE, Dec. 2016.

[12] R. Chignola and R. I. Foroni. Estimating the growth kinetics of experimental tumors from as few as two determinations of tumor size: implications for clinical oncology. IEEE transactions on biomedical engineering, 52(5):808–815, 2005.

[13] R. Chisholm, T. Lorenzi, A. Lorz, A. Larsen, L. Neves de Almedia, A. Escargueil, and J. Clairambault. Emergence of drug tolerance in cancer cell populations: an evolutionary outcome of selection, nongenetic instability, and stress-induced adaptation. Cancer Research, 75:930–939, 2015.

[14] A. Coldman and J. Goldie. A stochastic model for the origin and treatment of tumors containing drug-resistant cells. Bull. Math. Biol., 48:279–292, 1986.

[15] A. Correia and M. Bissell. The tumor microenvironment is a dominant force in multidrug resistance. Drug Resistance Updates, (15):39–49, 2012.

[16] J. Cunningham, R. Gatenby, and J. Brown. Evolutionary dynamics in cancer therapy. Mol. Pharm., 8:2094–2100, 2011.

[17] M. Dean, T. Fojo, and S. Bates. Tumour stem cells and drug resistance. Nature Reviews Cancer, (5):275–284, 2005.

[18] Y. E. Erdi. Limits of tumor detectability in nuclear medicine and pet. Molecular imaging and radionuclide therapy, 21(1):23, 2012.

[19] M. Feizabadi. Modeling multi-mutation and drug resistance: analysis of some cases. Theoretical Biology and Medical Modelling, 14:6, 2017.

[20] G. J. Fetterly, U. Aras, D. Lal, M. Murphy, P. D. Meholick, and E. S. Wang. Development of a preclinical pk/pd model to assess antitumor response of a sequential aflibercept and doxorubicin-dosing strategy in acute myeloid leukemia. The AAPS journal, 15(3):662–673, 2013.

[21] J. Foo, K. Leder, and S. Mumenthaler. Cancer as a moving target: understanding the composition and rebound growth kinetics of recurrent tumors. Evolutionary Applications, 6:54–69, 2013.

[22] J. Foo and F. Michor. Evolution of resistance to targeted anti-cancer therapies during continuous and pulsed administration strategies. PLoS Comput. Biol., 5:e1000557, 2009.

[23] J. Foo and F. Michor. Evolution of resistance to anti-cancer therapy during general dosing schedules. J. Theor. Biol., 263:179–188, 2010.

[24] J. Foo and F. Michor. Evolution of acquired resistance to anti-cancer therapy. J. Theor. Biol., 355:10–20, 2014.

[25] F. Fu, M. Nowak, and S. Bonhoeffer. Spatial heterogeneity in drug concentrations can facilitate the emergence of resistance to cancer therapy. PLoS Computational Biology, 11:e1004142, 2015.

[26] T. Gajewski, M. Y., C. Blank, I. Brown, K. A., J. Kline, and H. H. Immune resistance orchestrated by the tumor microenvironment. Immunological Reviews, (213):131–145, 2006.

[27] R. A. Gatenby, A. S. Silva, R. J. Gillies, and B. R. Frieden. Adaptive therapy. Cancer research, 69(11):4894–4903, 2009.

[28] J. Gevertz, Z. Aminzare, K.-A. Norton, J. Perez-Velazquez, A. Volkening, and K. Rejniak. Emergence of anti-cancer drug resistance: exploring the importance of the microenviron-mental niche via a spatial model. In T. Jackson and A. Radunskaya, editors, Applications of Dynamical Systems in Biology and Medicine, volume 158 of The IMA Volumes in Mathematics and its Applications, pages 1–34. Springer-Verlag, 2015.

[29] A. Goldman, B. Majumder, A. Dhawan, S. Ravi, D. Goldman, M. Kohandel, P. Majumder, and S. Sengupta. Temporally sequenced anticancer drugs overcome adaptive resistance by targeting a vulnerable chemotherapy-induced phenotypic transition. Nature Communications, 6:6139, 2015.

[30] M. Gottesman. Mechanisms of cancer drug resistance. Annual Review of Medicine, (531):615–627, 2002.

[31] J. Greene, O. Lavi, M. Gottesman, and D. Levy. The impact of cell density and mutations in a model of multidrug resistance in solid tumors. Bull. Math. Biol., 76:627–653, 2014.

[32] M. Hadjiandreou and G. Mitsis. Mathematical modeling of tumor growth, drug-resistance, toxicity, and optimal therapy design. IEEE Trans. Biomed. Eng., 61:415–425, 2013.

[33] C. Holohan, S. Van Schaeybroeck, D. B. Longley, and P. G. Johnston. Cancer drug resistance: an evolving paradigm. Nature Reviews Cancer, 13(10):714–726, 2013.

[34] G. Housman, S. Byler, S. Heerboth, K. Lapinska, M. Longacre, N. Snyder, and S. Sarkar. Drug resistance in cancer: an overview. Cancers, 6:1769–1792, 2014.

[35] Z. Iqbal, A. Aleem, M. Iqbal, M. Naqvi, A. Gill, and A. e. a. Taj. Sensitive detection of pre-existing BCR-ABL kinase domain mutations in CD34+ cells of newly diagnosed chronic-phase chronic myeloid leukemia patients is associated with imatinib resistance: implications in the post-imatinib era. PloS ONE, 8:e55717, 2013.

[36] T. Jackson and H. Byrne. A mathematical model to study the effects of drug resistance and vasculature on the response of solid tumors to chemotherapy. Mathematical Biosciences, 164:17–38, 2000.

[37] I. Koh, T. Hinoi, K. Sentani, E. Hirata, S. Nosaka, H. Niitsu, M. Miguchi, T. Adachi, W. Yasui, H. Ohdan, et al. Regulation of multidrug resistance 1 expression by cdx2 in ovarian mucinous adenocarcinoma. Cancer medicine, 5(7):1546–1555, 2016.

[38] N. Komarova and D. Wodarz. Drug resistance in cancer: Principles of emergence and prevention. Proc. Natl. Acad. Sci., 102:9714–9719, 2005.

[39] N. Komarova and D. Wodarz. Stochastic modeling of cellular colonies with quiescence: An application to drug resistance in cancer. Theor. Popul. Biol., 72:523–538, 2007.

[40] V. Krutyakov. Eukaryotic error-prone DNA polymerases: The presumed roles in replication, repair, and mutagenesis. Molecular Biology, 40(1):1–8, 2006.

[41] S. S. Lange, K.-i. Takata, and R. D. Wood. DNA polymerases and cancer. Nature Reviews Cancer, 11(2):96–110, 2011.

[42] O. Lavi, M. Gottesman, and D. Levy. The dynamics of drug resistance: A mathematical perspective. Drug Resist. Update, 15:90–97, 2012.

[43] O. Lavi, J. Greene, D. Levy, and M. Gottesman. The role of cell density and intratumoral heterogeneity in multidrug resistance. Cancer Res., 73:7168–7175, 2013.

[44] K. Leder, J. Foo, B. Skaggs, M. Gorre, C. Sawyers, and F. Michor. Fitness corrected by BCR-ABL kinase domain mutations determines the risk of pre-existing resistance in chronic myeloid leukemia. PLoS ONE, 6:e27682, 2011.

[45] U. Ledzewicz, K. Bratton, and H. Schättler. A 3-compartment model for chemotherapy of heterogeneous tumor populations. Acta Applicandae Mathematicae, 135:191–207, 2015.

[46] U. Ledzewicz and H. Schättler. Drug resistance in cancer chemotherapy as an optimal control problem. Discrete and Continuous Dynamical Systems - Series B, 5:129–150, 2006.

[47] W.-P. Lee. The role of reduced growth rate in the development of drug resistance of hob1 lymphoma cells to vincristine. Cancer letters, 73(2–3):105–111, 1993.

[48] L. Liu, F. Li, W. Pao, and F. Michor. Dose-dependent mutation rates determine optimum erlotinib dosing strategies for EGRF mutant non-small lung cancer patients. PLoS ONE, 10:e0141665, 2015.

[49] L. A. Loeb, C. F. Springgate, and N. Battula. Errors in dna replication as a basis of malignant changes. Cancer research, 34(9):2311–2321, 1974.

[50] A. Lorz, T. Lorenzi, J. Clairambault, A. Escargueil, and B. Perthame. Modeling effects of space structure and combination therapies on phenotypic heterogeneity and drug resistance in solid tumors. Bull. Math. Biol., 77:1–22, 2015.

[51] A. Lorz, T. Lorenzi, M. Hochberg, J. Clairambault, and B. Perthame. Populational adaptive evolution, chemotherapeutic resistance and multiple anti-cancer therapies. ESAIM: Math. Model. Num. Anal., 47:377–399, 2013.

[52] S. Luria and M. Delbrück. Mutations of bacteria from virus sensitivity to virus resistance. Genetics, 28:491–511, 1943.

[53] N. M. Makridakis and J. K. Reichardt. Translesion DNA polymerases and cancer. Frontiers in genetics, 3, 2012.

[54] D. McMillin, J. Negri, and C. Mitsiades. The role of tumour-stromal interactions in modifying drug response: challenges and opportunities. Nature Reviews Drug Discovery, (12):217–228, 2013.

[55] M. Meads, R. Gatenby, and W. Dalton. Environment-mediated drug resistance: a major contributor to minimal residual disease. Nature Reviews Cancer, (9):665–674, 2009.

[56] S. Menchón. The effect of intrinsic and acquired resistances on chemotherapy effectiveness. Acta Biother., 63:113–127, 2015.

[57] F. Michor, M. Nowak, and Y. Iwasa. Evolution of resistance to cancer therapy. Current Pharmaceutical Design, 12:261–271, 2006.

[58] S. Mumenthaler, J. Foo, N. Choi, N. Heise, K. Leder, D. Agus, W. Pao, F. Michor, and P. Mallick. The impact of microenvironmental heterogeneity on the evolution of drug resistance in cancer cells. Cancer Informatics, 14:19–31, 2015.

[59] S. Mumenthaler, J. Foo, K. Leder, N. Choi, D. Agus, W. Pao, P. Mallick, and F. Michor. Evolutionary modeling of combination treatment strategies to overcome resistance to tyrosine kinase inhibitors in non-small cell lung cancer. Mol. Pharm., 8:2069–2079, 2011.

[60] J. Nyce. Drug-induced DNA hypermethylation: a potential mediator of acquired drug resistance during cancer chemotherapy. Mutation Research, 386:153–161, 1997.

[61] J. Nyce, S. Leonard, D. Canupp, S. Schulz, and S. Wong. Epigenetic mechanisms of drug resistance: drug-induced DNA hypermethylation and drug resistance. Proceedings of the National Academy of Sciences, 90:2960–2964, 1993.

[62] P. Orlando, R. Gatenby, and J. Brown. Cancer treatment as a game: integrating evolutionary game theory into the optimal control of chemotherapy. Phys. Biol., 9:065007, 2012.

[63] J. Perez-Velazquez, J. Gevertz, A. Karolak, and K. Rejniak. Microenvironmental niches and sanctuaries: A route to acquired resistance. In K. Rejniak, editor, Systems Biology of Tumor Microenvironment, volume 936 of Advances in Experimental Medicine and Biology, pages 149–164. Springer, Cham, 2016.

[64] A. Pisco and S. Huang. Non-genetic cancer cell plasticity and therapy-induced stemness in tumour relapse: ‘What does not kill me strengthens me’. British Journal of Cancer, 112:1725–1732, 2015.

[65] A. O. Pisco, A. Brock, J. Zhou, A. Moor, M. Mojtahedi, D. Jackson, and S. Huang. Non-darwinian dynamics in therapy-induced cancer drug resistance. Nature communications, 4, 2013.

[66] C. Roche-Lestienne and C. Preudhomme. Mutations in the ABL kinase domain pre-exist the onset of imatinib treatment. Semin. Hematol., 40:80–82, 2003.

[67] S. E. Shackney, G. W. McCormack, and G. J. Cuchural. Growth rate patterns of solid tumors and their relation to responsiveness to therapy: an analytical review. Annals of internal medicine, 89(1):107–121, 1978.

[68] S. M. Shaffer, M. C. Dunagin, S. R. Torborg, E. A. Torre, B. Emert, C. Krepler, M. Beqiri, K. Sproesser, P. A. Brafford, M. Xiao, et al. Rare cell variability and drug-induced reprogramming as a mode of cancer drug resistance. Nature, 546(7658):431, 2017.

[69] A. Shah, K. Rejniak, and J. Gevertz. Limiting the development of anti-cancer drug resistance in a spatial model of micrometastases. Mathematical Biosciences and Engineering, (13):1185–1206, 2016.

[70] S. Sharma, D. Lee, B. Li, and M. e. a. Quinlan. A chromatin-mediated reversible drug-tolerant state in cancer cell subpopulations. Cell, 141:69–80, 2010.

[71] A. Silva and R. Gatenby. A theoretical quantitative model for evolution of cancer chemotherapy resistance. Biology Direct, 5:25, 2010.

[72] E. D. Sontag. Dynamic compensation, parameter identifiability, and equivariances. PLoS computational biology, 13(4):e1005447, 2017.

[73] L. H. Swift and R. M. Golsteyn. Genotoxic anti-cancer agents and their relationship to dna damage, mitosis, and checkpoint adaptation in proliferating cancer cells. International journal of molecular sciences, 15(3):3403–3431, 2014.

[74] B. Szikriszt, A. Póti, O. Pipek, M. Krzystanek, and N. e. a. Kanu. A comprehensive survey of the mutagenic impact of common cancer cytotoxics. Genome Biology, 17:99, 2016.

[75] H. Thomas, H. M. Coley, et al. Overcoming multidrug resistance in cancer: an update on the clinical strategy of inhibiting p-glycoprotein. Cancer control, 10(2):159–159, 2003.

[76] T. A. Traina and L. Norton. Log-kill hypothesis. In Encyclopedia of Cancer, pages 2074–2075. Springer, 2011.

[77] O. Trédan, C. Galmarini, and I. Tannock. Drug resistance and the solid tumor microenvironment. Journal of the National Cancer Institute, (99):1441–1454, 2007.

[78] B. Waclaw, I. Bozic, M. Pittman, R. Hruban, B. Vogelstein, and M. Nowak. A spatial model predicts that dispersal and cell turnover limit intratumor heterogeneity. Nature, 525:261–264, 2015.

[79] D. Woods and J. Turchi. Chemotherapy induced DNA damage response. Cancer Biology & Therapy, (14):379–389, 2013.

[80] Q. Wu, M.-y. Li, H.-q. Li, C.-h. Deng, L. Li, T.-y. Zhou, and W. Lu. Pharmacokineticpharmacodynamic modeling of the anticancer effect of erlotinib in a human non-small cell lung cancer xenograft mouse model. Acta Pharmacologica Sinica, 34(11):1427–1436, 2013.

[81] K. Yamamoto, K. Hirota, S. Takeda, and H. Haeno. Evolution of pre-existing versus acquired resistance to platinum drugs and PARP inhibitors BRCA-associated cancers. PLoS ONE, 9:e105724, 2014.

[82] H. Zahreddine and K. Borden. Mechanisms and insights into drug resistance in cancer. Frontiers in Pharmacology, (4):28, 2013.

